# The kinesin Kif21b regulates radial migration of cortical projection neurons through a non-canonical function on actin cytoskeleton

**DOI:** 10.1101/2023.02.06.526840

**Authors:** José Rivera Alvarez, Laure Asselin, Peggy Tilly, Roxane Benoit, Claire Batisse, Ludovic Richert, Julien Batisse, Bastien Morlet, Florian Levet, Noémie Schwaller, Yves Mély, Marc Ruff, Anne-Cécile Reymann, Juliette D. Godin

## Abstract

Completion of neuronal migration is critical for brain development. Kif21b is a plus-end directed kinesin motor protein that promotes intracellular transport and controls microtubule dynamics in neurons. Here we report a physiological function of Kif21b during radial migration of projection neurons in the mouse developing cortex. *In vivo* analysis in mouse and live imaging on cultured slices demonstrate that Kif21b regulates the radial glia-guided locomotion of new-born neurons independently of its motility on microtubules. Unexpectedly we show that Kif21b directly binds and regulates the actin cytoskeleton both *in vitro* and *in vivo* in migratory neurons. We establish that Kif21b-mediated regulation of actin cytoskeleton dynamics influences branching and nucleokinesis during neuronal locomotion. Altogether, our results reveal atypical roles of Kif21b on the actin cytoskeleton during migration of cortical projection neurons.

## Introduction

The cerebral cortex is a central structure of the mammalian brain that commands all higher-order functions. It contains a large number of specialized types of excitatory projection neurons, inhibitory interneurons and glia that are distributed within layers and are regionally organized into specialized areas responsible of motor, sensory, and cognitive functions (Greig et al., 2013; Jabaudon, 2017; Rash and Grove, 2006). Proper functioning of the neocortex depends on the active migration of the two major classes of cortical neurons, the pyramidal projection neurons that primarily engage in radial migration (Molyneaux et al., 2007) and the GABAergic interneurons that undergo tangential migration (Wonders and Anderson, 2006). Consequently, in human, abnormal cortical layering leads to cortical malformation and is often associated with epilepsy and intellectual disability (Barkovich et al., 2012; Fernandez et al., 2016; Francis and Cappello, 2020; Guerrini and Dobyns, 2014).

To reach their final position in the cortical plate, projection neurons first undergo somal translocation and then, as the cortical wall thickens, they switch to a multimodal radial migration that is accompanied by a series of highly coordinated morphological changes (Nadarajah et al., 2001; Noctor et al., 2004). Newborn projection neurons first adopt a multipolar morphology and migrate randomly in the intermediate zone (LoTurco and Bai, 2006; Tabata and Nakajima, 2003). Then, they undergo a multipolar-to-bipolar transition to initiate locomotion along the radial glia scaffold. During this locomotion phase, the displacement of projection neurons is paced by successive cycles of nucleokinesis (Martinez-Garay et al., 2016; Tanaka et al., 2004a; Tanaka et al., 2004b) as well as occasional pauses (Hurni et al., 2017). Once the glia-guided locomotion is completed, the projection neurons connect to the pia and undergo a final somal translocation to settle at appropriate position in the cortical plate (Nadarajah et al., 2001). Most of the identified mechanisms underlying the different steps of radial migration ultimately converge on the dynamic remodeling of both microtubule (MT) and actin cytoskeletons (Liaci et al., 2021; Lian and Sheen, 2015; Moon and Wynshaw-Boris, 2013; Wu et al., 2014). Further reflecting the importance of these cytoskeletons in the regulation of radial migration, variants in genes encoding for tubulin subunits, actin components, microtubule associated proteins, actin binding proteins or motor proteins have been largely associated to neuronal migration disorders (Francis and Cappello, 2020; Stouffer et al., 2016).

Kif21b is a kinesin particularly enriched in the brain from early development onwards (Asselin et al., 2020; Labonte et al., 2014; Marszalek et al., 1999), that promotes intracellular transport along microtubules (Ghiretti et al., 2016; Gromova et al., 2018; Muhia et al., 2016; van Riel et al., 2017) and controls MTs dynamics (Ghiretti et al., 2016; Hooikaas et al., 2020; Muhia et al., 2016; Taguchi et al., 2022; van Riel et al., 2017). In particular, Kif21b accumulates at the MT plus ends, where it stabilizes assembled MT by limiting MT polymerization and suppressing catastrophes (Hooikaas et al., 2020; Masucci et al., 2022; Taguchi et al., 2022; van Riel et al., 2017). During cortical development, Kif21b is specifically expressed in postmitotic neurons where it localizes in dendrites, axons and growth cones (Asselin et al., 2020; Huang and Banker, 2012; Marszalek et al., 1999). Depletion of Kif21b *in vivo* in mice leads to severe brain malformations including microcephaly, hydrocephaly and dysgenesis of the corpus callosum (Kannan et al., 2017), as well as deficits in learning, memory and social behavior (Gromova et al., 2018; Morikawa et al., 2018; Muhia et al., 2016) and impaired neuronal maturation and synaptic function (Morikawa et al., 2018; Muhia et al., 2016; Swarnkar et al., 2018). Although these results clearly point to critical roles of Kif21b in brain development and function, the underlying cellular and molecular mechanisms have not been elicited yet.

Pathological variants in *KIF21B* have been identified in patients presenting with brain malformation and intellectual disability (Asselin et al., 2020; Narayanan et al., 2022). Interestingly, those variants impair neuronal migration and interhemispheric connectivity by enhancing KIF21B canonical motor activity through a gain-of-function mechanism (Asselin et al., 2020; Narayanan et al., 2022). In addition, haploinsufficient *KIF21B* variant also leads to migratory defects, suggesting that dosage of KIF21B is critical for neuronal migration (Asselin et al., 2020), the exact underlying molecular mechanisms of which remains elusive.

Here we uncover the physiological roles of Kif21b during radial migration of cortical projection neurons. By combining *in vivo* genetic perturbation in mouse cortices, complementation assays and time-lapse recording on organotypic brain slices, we show that the kinesin Kif21b is required to maintain bipolar morphology and to promote nucleokinesis and dynamic branching of cortical projection neurons during glia-guided locomotion. Surprisingly, the canonical motor activity of Kif21b is dispensable for its function in migrating neurons. We demonstrate that Kif21b controls radial migration partly by regulating actin cytoskeleton dynamics through its direct binding to actin filaments. Altogether our data identify an unexpected non-canonical function of Kif21b on actin cytoskeleton participating to the regulation of cortical neurons migration.

## Results

### Kif21b is cell-autonomously required for radial migration of projection neurons

To explore the function of Kif21b on neuronal migration, we assessed the consequences of acute depletion of *Kif21b* specifically in postmitotic neurons using *in utero* electroporation (IUE) of CRE-dependent inducible shRNA vectors (Matsuda and Cepko, 2007) together with two plasmids allowing expression of CRE or GFP under the control of the NeuroD promoter (NeuroD:CRE and NeuroD:GFP respectively) in mouse embryonic cortices at E14.5. Efficacy of the two shRNAs was confirmed by Western blot in HEK-296T cell line (-93.4% for sh-*Kif21b* #1, -61.4% for sh-*Kif21b* #2 when compared to Kif21b expression in Scramble transfected cells) (**Figures S1A and S1B**). As described previously (Asselin et al., 2020), four days after IUE, the distribution of GFP-positive (GFP+) neurons depleted for *Kif21b* was significantly impaired with a notable reduction of GFP+ neurons reaching the upper cortical plate (Up CP) upon acute depletion of *Kif21b* compared to control (Scramble shRNA) (-19.4% for sh-*Kif21b* #1, -30% for sh-*Kif21b* #2) (**Figures 1A** and **1B**, **Figures S1C and S1D**). *Kif21b*-depleted projection neurons arrested in the intermediate zone expressed the upper-layer marker Cux1, supporting a faulty neuronal migration rather than specification defects (**Figure 1C**). As Kif21a, the other member of kinesin-4 family, shares a high sequence similarity with Kif21b (Marszalek et al., 1999), we tested whether the two paralogues have redundant function in neuronal migration. We assessed the ability of mouse wild-type (WT) Kif21b or Kif21a proteins to restore migration defects induced by *Kif21b* knock-down. We performed co-electroporation of plasmids expressing Kif21b or Kif21a under the regulation of the neuronal NeuroD promotor together with NeuroD-driven sh-*Kif21b* #2 that targets the 3’UTR of endogenous *Kif21b* transcripts (**Figure S1B**). Whereas Kif21b fully restored the defective migration induced by sh-*Kif21b* #2 (**Figures 1A and 1B, Figures S3B and S3C**), co-electroporation of Kif21a failed to rescue the impaired distribution of *Kif21b*-depleted neurons, suggesting that loss of Kif21b function in migrating neurons cannot be compensated by Kif21a. Notably, Kif21b likely regulated neuronal migration in a cell-autonomous manner as *Kif21b* silencing did not affect cell survival (**Figure S1E**) and and glia scaffold integrity (data not shown). Finally, we investigated the migration phenotype in *Kif21b^flox/flox^* conditional knock-out (cKO) embryos by IUE of NeuroD:Cre and NeuroD:GFP plasmids at E14.5. We confirmed the impaired positioning of *Kif21b* knock-out neurons four days after IUE, with a reduction of 13.9% of neurons distributed in the upper cortical plate in *Kif21b^flox/flox^* compared to *Kif21b^WT/WT^* embryos (**Figures 1D and 1E**).

**Figure 1.**
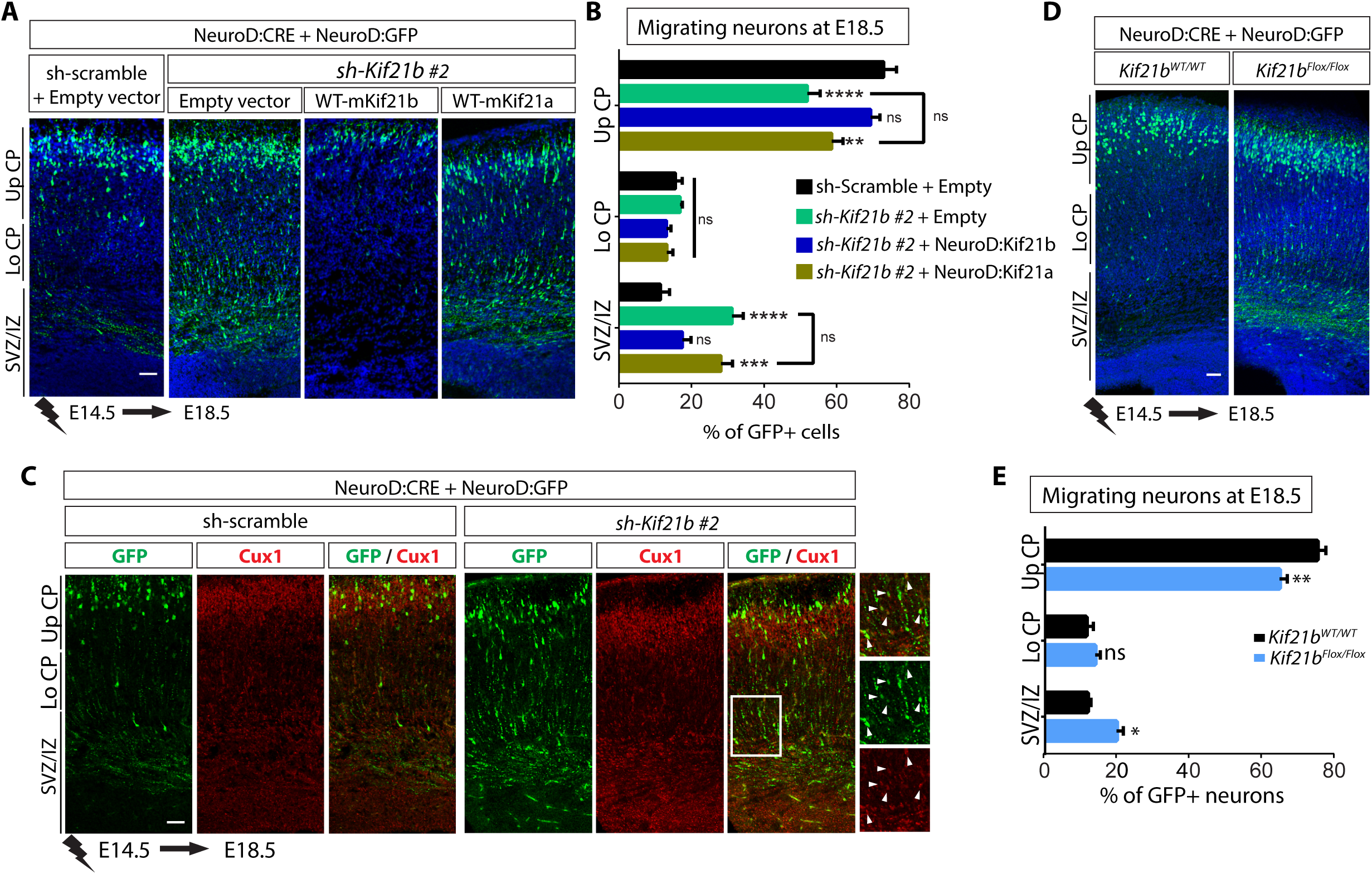
Kif21b is cell-autonomously required for radial migration of projection neurons. (**A**) Coronal sections of E18.5 mouse cortices electroporated at E14.5 with NeuroD:Cre and NeuroD:IRES-GFP together with either Cre inducible shRNA-*Kif21b* #2 in combination with NeuroD-Empty vector, NeuroD-mKif21b (WT(Wild-type)-mKif21b) or NeuroD-mKif21a (WT(Wild-type)-mKif21a) or sh-scramble in combination with NeuroD-Empty vector. (**D**) Coronal sections of E18.5 WT or double floxed *Kif21b* conditional knock-out (*Kif21b*^flox/flox^) mouse cortices electroporated at E14.5 with NeuroD:Cre and NeuroD:IRES-GFP. (**A, D**) GFP-positive electroporated cells are depicted in green. Nuclei are stained with DAPI. (**B, E**) Histograms (means ± s.e.m.) showing the distribution of GFP-positive neurons in different regions (Up CP, Upper cortical plate; Lo CP, Lower cortical plate; SVZ/ IZ, subventricular zone / intermediate zone) showing specific roles of Kif21b and not its paralogue Kif21a in projection neurons migration. Significance was calculated by two-way ANOVA, Bonferroni’s multiple comparisons test. ns, non-significant; *P < 0.05; **P < 0.005; ***P < 0.001, ****P < 0.0001. Number of embryos analyzed: (**B**) sh-scramble + Empty vector, n=6; sh-RNA-*Kif21b* #2 + Empty vector, n=11; sh-RNA-*Kif21b* #2 + WT-Kif21b, n=7; and sh-RNA-*Kif21b* #2 + WT-Kif21a, n=5. (**D**) Wild type, n=4; and *Kif21b*^flox/flox^, n=5. (**C**) Cux1 immunostaining (red) of E18 mouse cortices electroporated at E14.5 with NeuroD:Cre and NeuroD:IRES-GFP together with either Cre inducible sh-scramble or shRNA *Kif21b* #2 showing correct specification of GFP-positive *Kif21b*-depleted arrested neurons (arrowheads in insets). Scale bars (**A**, **C**, **D**) 50 μm. See also Figures S1 and S2.

Given the expression of Kif21b in the mouse developing ganglionic eminences where the cortical inhibitory GABAergic neurons are born (**Figures S2A and S2B**), we next asked whether Kif21b also regulates tangential migration of interneurons. We performed *in vitro* culture of medial ganglionic explants (MGE) from E13.5 Dlx:Cre-GFP embryos electroporated *ex-vivo* with Cre dependent cherry expressing vector (pCAG-loxPSTOPloxP-Cherry) together with CRE-inducible sh-scramble or sh-*Kif21b* #2 on a layer of WT cortical feeders. This strategy allows concomitant knockdown of *Kif21b* and expression of a cherry fluorescent reporter specifically in migrating cortical interneurons. Time lapse recordings of electroporated explants revealed that, although *Kif21b*-silenced interneurons were able to migrate out of the explant, their kinetics of migration were different from the sh-scramble electroporated interneurons (**Figures S2C and S2D**), with a decreased migration velocity of 24.3%. Altogether, our results revealed the kinesin Kif21b as a key regulator of neuronal migration of the two main classes of cortical neurons.

### Kif21b depletion impedes glia-guided locomotion of projection neurons

To determine how Kif21b controls migration of cortical projection neurons, we further investigated the dynamic cell shape remodeling occurring at the successive steps of migration along radial glia fibers (**Figure 2A**) by time lapse recording. *In utero* electroporation of inducible *Kif21b* shRNA or sh-scramble together with NeuroD:Cre and NeuroD:GFP plasmids were performed in WT E14.5 embryos. Two days later brain organotypic slices were cultured in vitro and GFP+ neurons were recorded the next day for 10 hours (**Figures 2B and 2C**). Although *Kif21*b depletion did not affect the initial multipolar to bipolar transition required to start glia-guided locomotion (**Figure 2A**, step 1; **Figure 2D**), the percentage of bipolar neurons that actively initiated the locomotory phase was decreased by 13.8% in sh-*Kif21b* #2 condition compared to sh-scramble control (76% and 88.1% respectively) (**Figure 2A**, step 2; **Figure 2E**). Moreover, compared to control neurons, the *Kif21b*-silenced bipolar neurons that started glia-guided locomotion (step 3 in **Figure 2A**) migrated significantly slower (**Figures 2B-C and 2F**) and spent more time pausing (**Figures 2C and 2G**). Exacerbated pausing was due to increases in both frequency (**Figure 2H**) and duration of the pauses (**Figure 2I**), likely due to a loss of bipolar polarity as shown by *Kif21b*-depleted neurons converting more often to multipolar shape while pausing (step 4 in **Figure 2A**, **Figure 2J**). Interestingly, the motility index, defined as the velocity during active locomotion excluding the pauses, was decreased in *Kif21b*-silenced neurons (step 5 in **Figure 2A, Figure 2K**), demonstrating that Kif21b regulates both pausing and effective processivity of migrating projection neurons.

**Figure 2.**
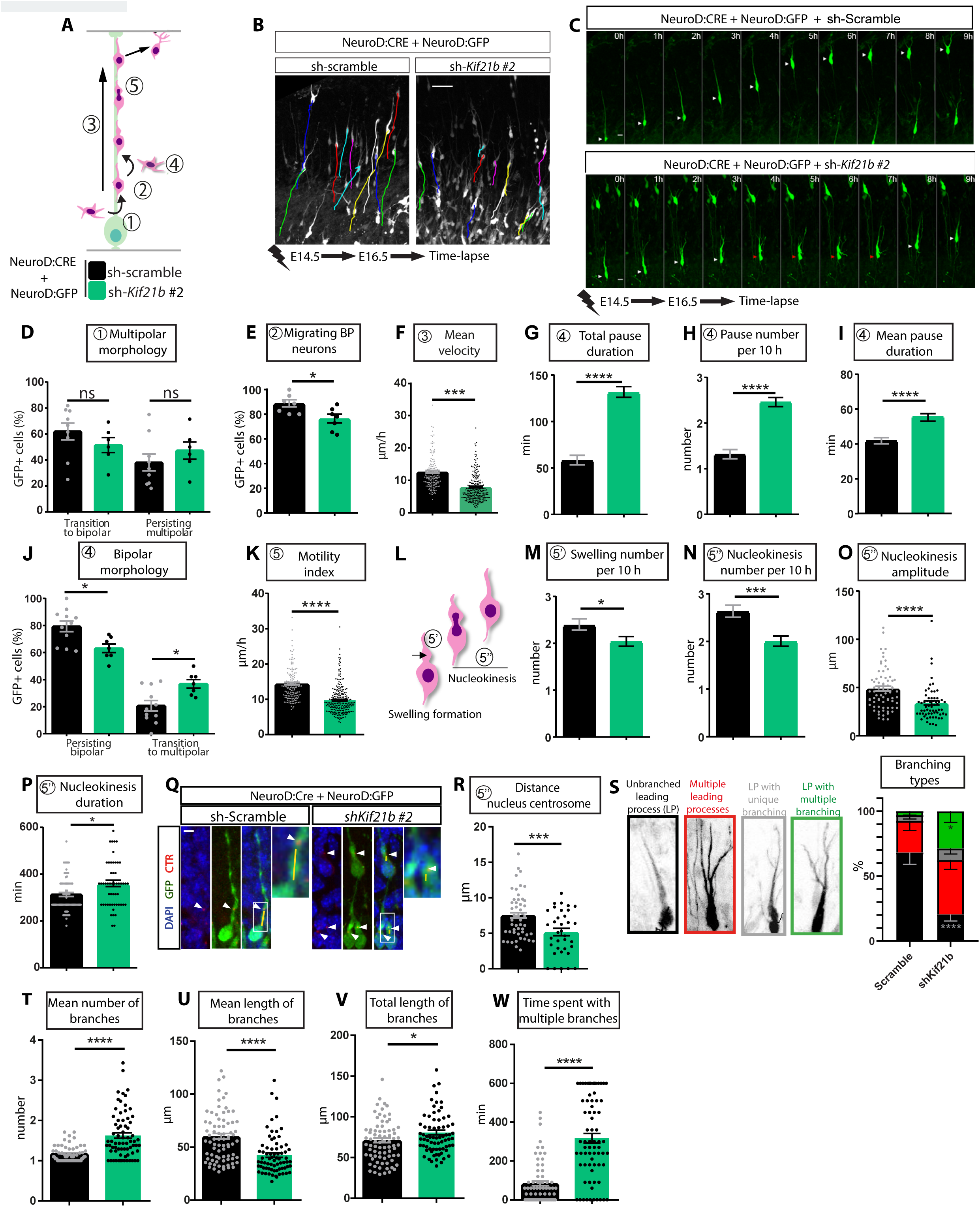
Kif21b depletion impedes glia-guided locomotion of projection neurons. (**A**) Drawing of radial migration of projection neurons depicting the different steps of the process, step 1: multipolar-bipolar transition, step 2: initiation of bipolar migration, step 3: glia-guided locomotion that is paced by pauses (step 4) when bipolarity is transiently loss and active locomotion (step 5). (**B-W**) Time lapse imaging of neurons electroporated at E14.5 with NeuroD:Cre and NeuroD:IRES-GFP and either Cre inducible shRNA *Kif21b* #2 or sh-scramble in E16.5 brain slices cultured for one day. (**B**) Locomotor paths (colored lines) of sh-scramble or sh-*Kif21b* #2 expressing neurons recorded every 30 minutes for 10 hours (h). Scale bar: 20 μm. (**C**) Time-lapse sequences of representative pyramidal neurons electroporated with the indicated constructs at E14.5. White and red arrowheads indicate respectively forward movement and pausing of the nucleus. Scale bar: 20 μm. (**D**) Percentage of GFP+ neurons converting to a bipolar morphology or remaining as multipolar for 10 hours recording. (**E-K**) Quantification (means ± s.e.m.) of the percentage of GFP+ bipolar neurons (BP) initiating locomotion (**E**), the mean velocity of locomotion (μm/h) (**F**),the total pause duration (min) (**G**), the average number of pauses per 10 hours recording (**H**), the mean pause duration (min) (**I**), the percentage of neurons maintaining bipolar shape or converting to multipolar morphology (**J**), and the motility index (velocity without pauses (μm/h)) (**K**). (**L**) Schematic representation of the forward progression of pyramidal neurons during glia guided locomotion, involving cycles of swelling formation (black arrow in 5’) and nucleokinesis (5’’). (**M-P**) Analysis (means ± s.e.m.) of the average number of swelling (**M**) or nucleokinesis (**N**) during 10 hours recording and mean amplitude (μm) (**O**) and duration (min) (**P**) of each nucleokinesis. (**Q**) Representative confocal images of migrating neurons after GFP immunolabelling (green) of E16.5 mouse cortices electroporated at E14.5 with NeuroD:Cre, NeuroD:GFP and pCAGGS-PACT-mKO1 to label centrosome (CTR, red, arrowheads), and either Cre inducible shRNA *Kif21b* #2 or sh-scramble. Nuclei were stained with DAPI (blue). Distances between nuclei and centrosome are indicated by a yellow line. Scale bar: 5 μm. (**R**) Quantification (means ± s.e.m.) of the distance nucleus-centrosome (μm) in neurons extending a swelling in control and *Kif21b-*depleted cells. (**S-W**) Analysis (means ± s.e.m.) of dynamic branching of the leading process showing branching types (**S**), number (**T**), mean length (μm) (**U**), total length (μm) (**V**) of branches, as well as time spent with multiple branches (min) (**W**). Significance was calculated by unpaired t-test, except in figure S, where significance was calculated by two-way ANOVA. ns, non-significant; *P < 0.05; ***P < 0.001, ****P < 0.0001. Number of embryos analyzed: (**D**) sh-scramble, n=9; sh-*Kif21b* #2, n=6; (**E**) sh-scramble, n=7; sh-*Kif21b* #2, n=7; (**J**) sh-scramble, n=11; sh-*Kif21b* #2, n=7. Number of cells analyzed from at least 3 embryos: (**F-I,K**) sh-scramble, n=153; sh-*Kif21b* #2, n=248; (**M, N**) sh-scramble, n=77; sh-*Kif21b* #2, n=83; (**O,P**) sh-scramble, n=72; sh-*Kif21b* #2, n=62; (**R**) sh-scramble, n=51; sh-*Kif21b* #2, n=35, (**S-W**) sh-scramble, n=73; sh-*Kif21b* #2, n=71.

To properly migrate, projection neurons rely on repetitive cycles of forward progression of the cell body that starts with the translocation of a cytoplasmic dilatation (also known as a swelling) in the proximal region of the extending leading process (Step 5’ in **Figure 2L**) and that is followed by the nuclear translocation (hereafter called nucleokinesis) toward the cytoplasmic swelling (Nishimura et al., 2017) (Step 5’’ in **Figure 2L**). We showed that *Kif21b*-silenced neurons form less swellings and undergo less frequent and less efficient nucleokinesis with a net nuclear movement of 32.9 μm and 47.8 μm in *kif21b*-depeted and control neurons respectively (**Figures 2M-2P**). Defective nucleokinesis was further confirmed by shorter maximal distances between the nucleus and the centrosome, that is encompassed in the swelling, in neurons electroporated with sh-*Kif21b* #2 (**Figures 2Q and 2R**). We then investigated the dynamic branching of migrating neurons during active locomotion excluding the phases of pausing. While most control neurons persist bipolar during the locomotory phase with a single leading process that branched very transiently, *Kif21b*-depleted neurons, although still polarized, were characterized by multiple and heavily branched leading processes (**Figures 2S**). Overall, sh-*Kif21b* #2 expressing neurons showed an increased total branch length that arose from an increased number of shorter branches sprouting from either the soma or the leading process (**Figures 2S-2V**). As a result, neurons depleted for *Kif21b* spent statistically more time with multiple branches (**Figure 2W**), likely contributing, together with impaired nucleokinesis, to the decrease in migration velocity index observed in mutant projection neurons (**Figure 2K**). Collectively our data indicate that the kinesin Kif21b is critical to regulate neuronal polarization, nucleokinesis and dynamic branching of projection neurons during glia-guided locomotion.

### Kif21b motility on microtubules is dispensable for its function in migrating projection neurons

Given the critical role of Kif21b in regulating intracellular transport in neurons (Ghiretti et al., 2016; Gromova et al., 2018; Muhia et al., 2016; van Riel et al., 2017), we assessed whether Kif21b processive activity on microtubules mediate Kif21b function in migrating neurons. First, we tested for restoration of the sh-Kif21b #2-induced phenotype by truncated Kif21b protein that lacks the motor domain (Kif21bΔMD, **Figure 3A**) by introducing full-length or Kif21bΔMD (NeuroD:Kif21b) together with inducible sh-Kif21b #2 and NeuroD:CRE in projection neurons using *in utero* electroporation at E14.5. Interestingly, Kif21b protein that lacks motor domain only partially rescued the distribution of *Kif21b*-depleted neurons with a significant number of neurons still trapped in the intermediate zone at E18.5 (**Figures 3B and 3C**), suggesting that this domain is partially required for Kif21b function in migrating neurons. To further test for the need of Kif21b motility on microtubules for neuronal migration, we performed complementation assays by expressing Kif21b protein that either cannot bind (NeuroD:Kif21bΔATP; **Figure 3A**) or hydrolyze ATP (NeuroD:Kif21b-T96N; **Figure 3A**) (Muhia et al., 2016) in *Kif21b*-silenced neurons. Surprisingly, both constructs fully restored the migration phenotype at E18.5, demonstrating that motility of Kif21b is dispensable for its function during radial migration (**Figures 3B and 3C**). Accordingly, the carboxy-terminal WD40 domain of Kif21b (**Figure 3A**), that is thought to promote interaction with cargoes (Marszalek et al., 1999), is not essential for Kif21b migration regulatory activities, as radial migration phenotype is fully rescued by overexpression of Kif21bΔWD40 construct (**Figures 3B and 3C**). Of note, Kif21b truncated or mutant proteins showed similar expression level in N2A neuroblastoma cell line, except for NeuroD:Kif21bΔATP that tends to be lowly expressed (**Figure S3A**) and do not induce migration phenotypes while overexpressed under control conditions (Sh-Scramble) (**Figures S3B and S3C**). We next sought to investigate the migration steps that are regulated by the motor domain of Kif21b. We performed time lapse recording of brain organotypic slices after *in utero* electroporation of Kif21bΔMD construct together with inducible *Kif21b*-shRNA. In accordance with a partial rescue of *Kif21b*-silenced neurons distribution in the cortical plate, expression of Kif21b lacking the motor domain induced a rescue of 52.6% of the migration velocity of *Kif21b* depleted neurons (**Figures 3D and 3E**). This partial rescue is likely due to inability of Kif21bΔMD to restore pausing defects. Indeed, while Kif21bΔMD fully recovered the motility index phenotype (distance with respect to time minus pauses duration) (**Figure 3F**), it failed to restore the increased number of pauses (**Figure 3G**) and even induced more drastic defects on pauses duration (**Figures 3H and 3I**). In line, Kif21bΔMD expression did not rescue the loss of bipolar morphology of locomoting *Kif21b*-depeleted neurons (**Figure 3J**). Collectively those data indicate that the N-terminal motor domain of Kif21b independently of its motility on microtubules is critical to prevent pausing of of cortical projection neurons during migration.

**Figure 3.**
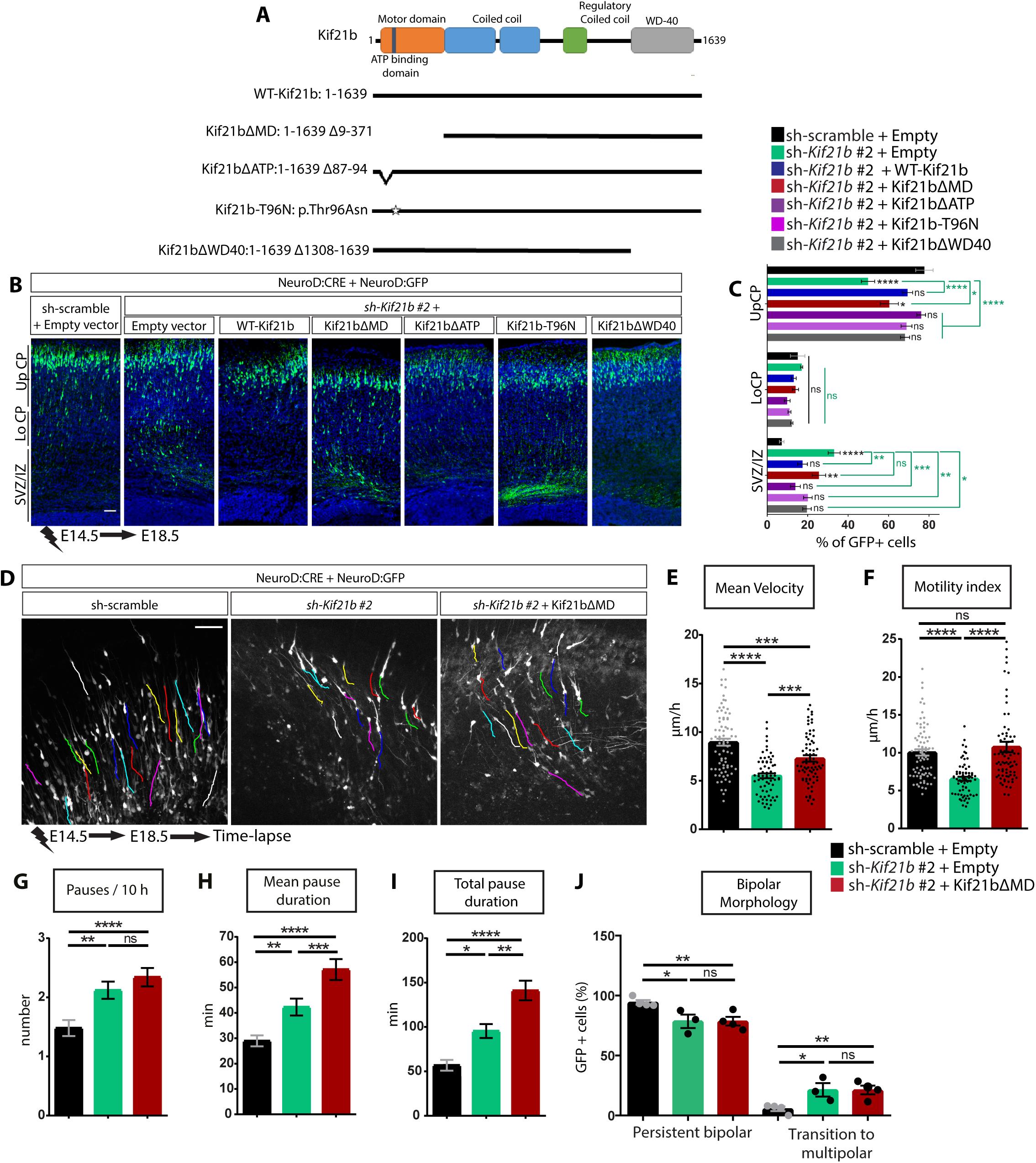
Kif21b motility on microtubules is not essential for its function in migrating projection neurons. (**A**) Schematic representation of the mouse Kif21b protein indicating its different domains, and the recombinants constructs used for the rescue experiments. (**B**) (**A**) Coronal sections of E18.5 mouse cortices electroporated at E14.5 with NeuroD:Cre and NeuroD:IRES-GFP together with either Cre inducible shRNA-*Kif21b* #2 in combination with NeuroD-Empty vector or Kif21b-expressing constructs as indicated, or sh-scramble in combination with NeuroD-Empty vector. GFP-positive electroporated cells are depicted in green. Nuclei are stained with DAPI. Scale bar, 50 μm. (**C**) Histograms (means ± s.e.m.) showing the distribution of GFP-positive neurons in different regions (Up CP, Upper cortical plate; Lo CP, Lower cortical plate; or SVZ /IZ, subventricular zone / intermediate zone) in all conditions as indicated. Number of embryos analyzed per condition: sh-scramble + Empty vector, n=3; sh-RNA-Kif21b + Empty vector, n=11; sh-Kif21b + indicated Kif21b constructs, n≥5. Significance was calculated by two-way ANOVA, Bonferroni’s multiple comparisons test. ns, non-significant; *P < 0.05; **P < 0.005; ***P < 0.001, ****P < 0.0001. (**D-J**) Time lapse imaging of neurons electroporated at E14.5 with NeuroD:Cre and NeuroD:IRES-GFP together with either Cre inducible shRNA-*Kif21b* #2 in combination with NeuroD-Empty vector or NeuroD:Kif21bΔMD, or sh-scramble in combination with NeuroD-Empty vector in E16.5 brain slices cultured for one day. (**D**) Locomotor paths (colored lines) of neurons expressing sh-scramble or sh-*Kif21b* #2 alone or in combination with NeuroD:Kif21bΔMD and recorded every 30 min for 10 hours (h). Scale bar: 20 μm. (**E-I**) Analysis (means ± s.e.m.) of migration dynamics over 10 hours showing that expression of NeuroD:Kif21bΔMD in Kif21b-depleted neurons partially and totally rescue the mean velocity (μm/h) (**E**) and the velocity index (μm/h) (**F**), respectively, but does not rescue the average number of pauses per 10 hours (**G**), the mean pause duration (min) (**H**), the total pause duration (min) (**I**) or the percentage of neurons converting to multipolar morphology (**J**). Significance was calculated by two-way ANOVA, Bonferroni’s multiple comparisons test. ns, non-significant; *P < 0.05; **P < 0.005; ***P < 0.001, ****P < 0.0001. Number of cells analyzed from at least 3 embryos: (**E-I**) sh-scramble + NeuroD-Empty, n=77; sh-*Kif21b* #2 + NeuroD-Empty, n=66, sh-*Kif21b* #2 + NeuroD:Kif21bΔMD, n=76. Number of embryos analyzed: (**J**) sh-scramble + NeuroD-Empty, n=4; sh-*Kif21b* #2+ NeuroD-Empty, n=3; sh-*Kif21b* #2+ NeuroD:Kif21bΔMD, n=4. See also Figure S3.

### Kif21b regulates actin cytoskeletal dynamics through direct binding

We next sought to understand the function of Kif21b that mediates its roles during radial migration of projection neurons. First, to probe for candidate proteins that may interact with Kif21b, we performed Kif21b co-immunoprecipitation on E18.5 mouse cortices followed by mass spectrometry analysis. Among the 41 proteins identified (**Table S1**), 12 were proteins associated either to the microtubule cytoskeleton or, more unexpectedly, to the actin cytoskeleton (**Figures 4A and S4A**). Among those cytoskeleton candidates, we found several actin paralogues, actin-based motor proteins including components of the non-muscle myosin II (NM2) motor protein, that powers actin contractile movement (Brito and Sousa, 2020), the unconventional myosin Va that transports cargoes along actin filaments (Hammer and Burkhardt, 2013) and the actin binding protein Drebrin (Dbn1). Given the predominant role of the actomyosin remodeling in neuronal cells (Schaar and McConnell, 2005; Solecki et al., 2009; Tsai et al., 2007) and the increasing evidences showing that disruption of actin cytoskeleton associates with neuronal migration defects in human (Francis and Cappello, 2020; Stouffer et al., 2016), the actin cytoskeleton emerged as a strong unlooked-for candidate to mediate Kif21b function in migrating neurons. To validate these interactions, we first performed anti-Kif21b immunoprecipitation assays on extracts from E18.5 wild-type cortices and confirmed the specific binding of Kif21b to both actin and myosin light chain 2 (MLC 2), a subunit of NM2 (**Figures 4B and 4C**). Reciprocal immunoprecipitation analysis using anti-actin antibody confirmed the specific interaction between Kif21b and actin (**Figure 4B**). Next, we purified actin from E18.5 cortices and detected Kif21b in the pellet containing polymerized filamentous actin (F-actin) (**Figure 4D**), further suggesting ability of Kif21b to interact with filaments of actin. Corroborating these findings, Kif21b is enriched in the actin-rich growth cone fraction isolated from postnatal mouse cortices (**Figure 4E**). Immunolabelling of Kif21b, actin and microtubules in primary cortical neurons at day *in vitro* 2 (DIV2) revealed localization of Kif21b both in axons and the central domain of the growth cone that are devoid of microtubules and essentially composed of actin networks (**Figure 4F**). These data suggest that neuronal Kif21b likely interacts with actin cytoskeleton independently of its binding to microtubules. To ascertain the localization of Kif21b on the actin cytoskeleton, we analyzed relative localization of endogenous Kif21b and the actin cytoskeleton labelled with phalloidin using multicolor Single Molecule Localization Microscopy (SMLM) in primary cortical neurons at DIV2. STORM imaging revealed that Kif21b molecules organized as clusters localized along actin filaments following a spiral pattern in the growth cone as well as in the neurites (**Figure 4G**). We performed quantitative analysis, based on the identification of objects from a Voronoi diagram (Levet et al., 2015), of the shortest distance between the centroid of each Kif21b cluster and the actin lattice border. Computation of the distances indicated that most Kif21b clusters localized within 1.34 nm and 1.86 nm (median) from the actin filament (**Figure 4H**) in growth cone and neurites respectively, demonstrating very close spatial approximation of Kif21b with actin filaments in cortical neurons. We next explored whether Kif21b directly binds to actin by performing *in vitro* co-sedimentation assays of pre-polymerized actin with purified recombinant Kif21b (**Figure S4B, Table S2**). Notably, although it contained negligible protein contaminants (**Table S2**), Kif21b purified fractions comprised full-length (FL) His-tagged Kif21b as well as shorter Kif21b fragments (**Table S2**). We detected FL Kif21b as well as most of the short products in the actin-enriched pellet (**Figure 4I**), suggesting a direct binding of Kif21b to polymerized actin. Accordingly, the same experiments in absence of actin revealed that most Kif21b fragments were found in the supernatant. As a control, we also performed a microtubule co-sedimentation assay with Kif21b purified fraction, showing expected enrichment of Kif21b in the MT pellet (data not shown). Finally, we investigated the roles of Kif21b on actin dynamics by recording, using spinning disc microcopy, dynamics of purified actin labelled with phalloidin and incubated with 1 nM of recombinant Kif21b. Actin polymerization was enhanced in presence of 1 nM of Kif21b (**Figure 4J**), indicating that direct binding of Kif21b may favor the initial step of actin dynamics, meaning nucleation (**Figure 4N**). Notably, increasing the concentration of Kif21b to 10 nM had reverse effect on actin nucleation (**Figure S4C**). Conversely, low concentration of Kif21b (1 nM) does not affect assembly of actin when actin is used at higher concentration (5 μM) (data not shown), suggesting that effect of Kif21b on actin nucleation depends on the relative stoichiometry of actin and Kif21b molecules. To validate this actin-related function in cortices, we performed spinning disc analysis of dynamics of mouse embryonic brain actin labelled with phalloidin using protein extracts from E18.5 WT and KO (*Kif21b*^tm1a/tm1a^) cortices. Corroborating assays with recombinant proteins, loss of Kif21b in embryonic cortices significantly limits *de novo* assembly of actin filaments as expected for a reduction of actin nucleation capacity (**Figure S4D**). Further analyzes of the G/F actin ratio by ultracentrifugation sedimentation assay in E18.5 WT and *Kif21b* KO (*Kif21b*^tm1a/tm1a^) cortices revealed a decreased proportion of the filamentous form of actin (F-actin) upon deletion of *Kif21b* compared to the control condition (**Figure 4K**), although measured total actin levels by Western Blot were increased in KO condition (**Figure 4L**). This confirms the positive role of Kif21b in regulating the nucleation of actin filaments in embryonic cortices (**Figure 4N**). Finally, tracking of individual actin filament revealed that Kif21b did not affect filament elongation as similar rates of actin filament growth were measured in presence of Kif21b and in actin alone condition (**Figure 4M**). Altogether these data converged towards a direct role of Kif21b in regulation of actin dynamics (**Figure 4N**).

**Figure 4.**
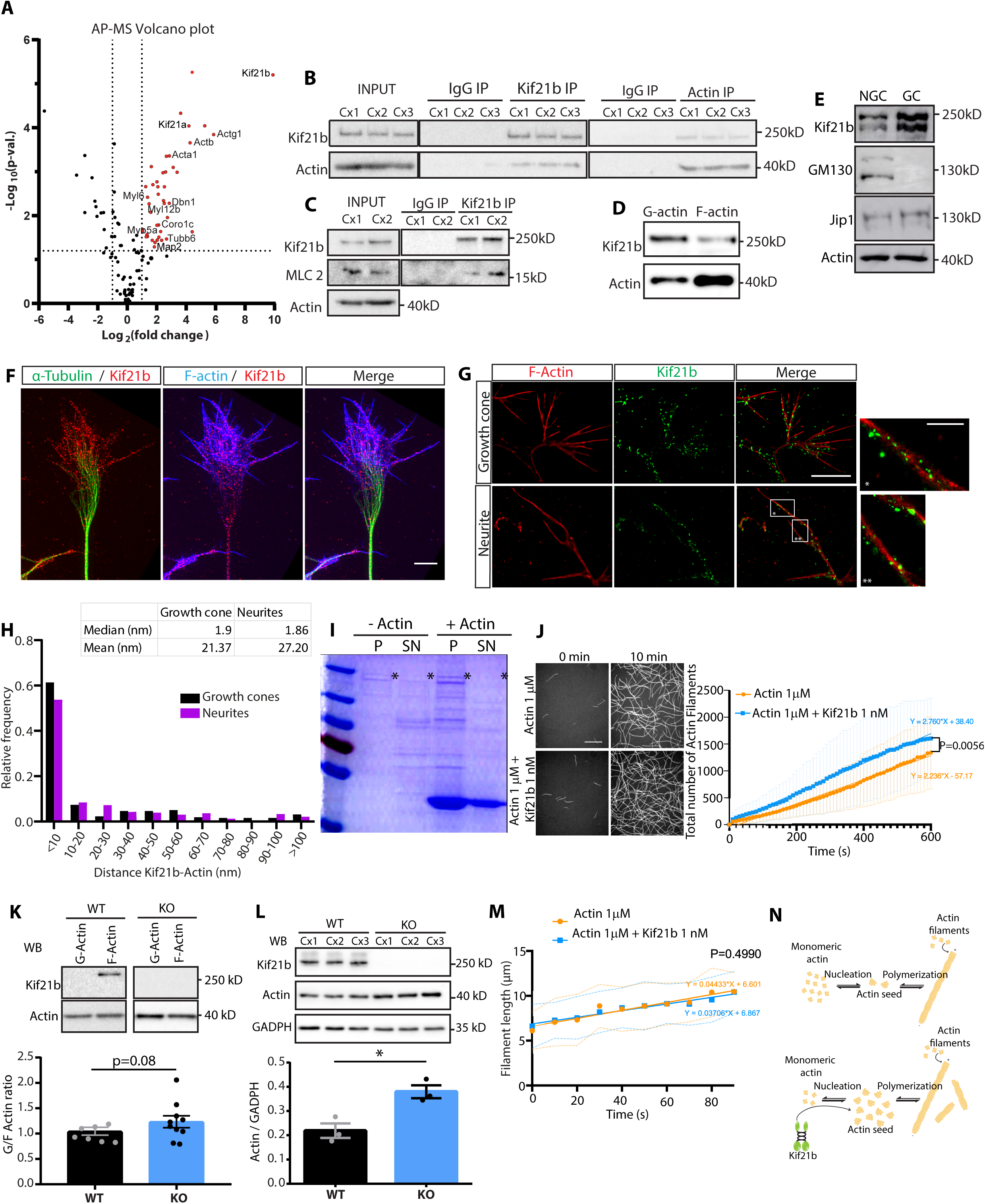
Kif21b regulates actin cytoskeletal dynamics through direct binding. (**A**) Volcano plot analysis of the affinity purification-mass spectrometry (AP-MS) results showing, in red, proteins that are specifically interacting with Kif21b in E18.5 mouse cortices. (**B-C**) Western blot analysis of Kif21b or actin immunoprecipitation (IP) experiments in 3 independent E18.5 mouse cortices (Cx 1 to 3) showing reciprocal interaction between Kif21b and actin (**B**) and binding of Myosin light chain-2 (MLC2) to Kif21b (**C**). Whole cortices extracts are shown as input. (**D**) Western blot F-actin sedimentation assay from E18.5 wild type mouse cortices showing that Kif21b binds to F-actin. (**E**) Growth cone fractionation assay showing growth cone (GC, enriched for vesicle marker Jip1) and a non-growth cone fraction (NGC, enriched for Golgi marker GM130) isolated from P2 wild type mouse cortices and indicating that Kif21b is enriched in the GC fraction. (**F**) Triple staining of Kif21b (red), α-tubulin (green) and F-actin (phalloidin, in blue) in a growth cone of a DIV2 neuron imaged by confocal microscopy. Scale bar: 5 μm. (**G**) Single Molecule Localization Microscopy (SMLM) showing Kif21b (green) and F-actin (phalloidin, red) double staining in a growth cone and a neurite of a DIV2 neuron. Scale bar: 5 μm. Insets are shown by one (*) or two asterisks (**). Scale bar in insets: 1 μm. (**H**) Analysis of the distribution probability of the distance (nm) between Kif21b clusters and actin filaments in both growth cones and neurites. Growth cones, n= 259 particles analyzed from 3 neurons, neurites, n= 331 particles analyzed from 4 neurons, experiment done in two independent replicates. (**I**) In vitro co-sedimentation assay of recombinant Kif21b in presence or absence of actin, showing that full-length Kif21b (asterisk) and shorter fragments co-sediment with polymerized actin found in the pellet (P). SN= supernatant. (**J**) Representative spinning-disk images of *in vitro* analysis of actin polymerization dynamics (stained with phalloidin) at the beginning (0’) and at the end (10’) of the experiment in presence or absence of recombinant Kif21b (1 nM). Scale bar 10 μm. The graph shows the number of actin filaments assembled during a 600 s time-lapse. Significance was calculated by linear regression to determine differences in curves for each dataset. Number of replicates: Actin 1 μM;, n=7, Actin 1 μM + Kif21b 1 nM, n= 6 in three different experiments. (**K**) Quantitative analysis (mean ± s.e.m.) of Western blot of actin sedimentation assay performed in E18.5 wild type (WT) and *Kif21b*^Tm1a/Tm1a^ (KO) mouse cortices showing increase of G/F ratio upon deletion of *Kif21b*. (**L**) Western blot analysis of total actin levels in three independent E18.5 wild type (WT) and *Kif21b*^Tm1a/Tm1a^ (KO) mouse cortices (Cx 1 to 3) showing (means ± s.e.m.) increased actin level normalized by GADPH in KO samples. (**K,L**) Significance was calculated by unpaired t-test, *P < 0.05. Number of embryos analyzed: (**K**) WT, n=7; KO, n=10; (**L**) WT and KO, n=3. (**M**) *In vitro* actin polymerization in absence or presence of recombinant Kif21b (1 nM) analyzed as the length of individual filaments of actin during a 80 s time-lapse. Significance was calculated by linear regression to determine differences in curves for each dataset. Number of filaments analyzed: Actin 1 μM, n=12, Actin 1 μM + Kif21b 1 nM, n= 14 in three different experiments. (**N**) Scheme representing monomer actin pool nucleation and polymerization. Lower panel showed the positive effect of Kif21b on actin nucleation without affecting polymerization. See also Figure S4.

### Kif21b depletion leads to aberrant actin dynamics in migrating neurons

Although well studied in the context of tangential migration of interneurons (Bellion et al., 2005; Godin et al., 2012) or glia-guided migration of cerebellar granular neurons (Solecki et al., 2009), the actin cytoskeletal dynamics in migrating cortical projection neurons has been poorly addressed. We first examined the spatial distribution of actin during glia-guided locomotion of cortical projection neurons using Lifeact as a probe for F-actin (Riedl et al., 2008) in brain organotypic slices after *in utero* electroporation of LifeAct-Ruby in E14.5 cortices. Time lapse imaging in control neurons (sh-Scramble) revealed that LifeAct-labelled F-actin accumulated in the proximal part of leading process before the translocation of the nucleus and dropped severely once the nuclear movement is completed (**Figures 5A and 5B**). Of note, in this experimental setup, the sensitivity of the LifeAct was not compatible with analysis of actin dynamics during branching. Interestingly, although actin dynamics were similar in control and *Kif21b*-depleted neurons before nucleokinesis, F-actin accumulation in the leading process persisted after the forward movement of the nucleus in neurons electroporated with sh-*Kif21b* #2 (**Figures 5A and 5C**). These data demonstrated the critical role of Kif21b in regulating actin dynamics *in vivo* in migrating neurons. Given the predominant role of actomyosin contraction in promoting nuclear translocation of migrating neurons (Schaar and McConnell, 2005; Solecki et al., 2009; Tsai et al., 2007) and the binding of Kif21b to non-muscle myosin II (**Figures 4B and 4C**), we tested the ability of Blebbistatin, which inhibits actomyosin contraction through inhibition of Myosin II motor activity, to rescue the dynamics of migrating neurons. Organotypic slices of brain electroporated with inducible sh-*Kif21b* #2, NeuroD:CRE and NeuroD:GFP were recorded for 4 hours, then treated with Blebbistatin at 3μM and imaged for additional 3.5 hours (**Figure 5D**). Nucleokinesis number and amplitude were similar before and after Blebbistatin treatment (**Figures 5E and 5F**), suggesting that faulty nucleokinesis induced by *Kif21b* depletion does not rely on elevated myosin activity and actomyosin contraction. As we demonstrated that Kif21b regulates migration pausing independently of its motility on microtubules (**Figures 3G-I**), we tested whether pausing could be related to actin-dependent Kif21b function. We showed that inhibition of actomyosin contraction by Blebbistatin failed to rescue number (**Figure 5G**) and duration of pauses (**Figures 5H-and 5I**) indicating that Kif21b likely regulates neuronal polarization independently of actomyosin contractility (**Figure 5J**). Finally, we showed that inhibition of Myosin II motor activity with Blebbistatin rescued the multiple branching observed in *Kif21b*-depleted neurons (**Figures 5K and 5L**), suggesting that Kif21b, at least partly, regulates dynamic branching of the leading process through actomyosin dependent mechanisms. However, the partial rescue of the time spent with multiple branches (26min/3.5h in control condition (**Figure 2W**), 94min/3.5h sh-*Kif21b* #2 pre-Blebbistatin, 41min/3.5h sh-*Kif21b* #2 post-Blebbistatin) is likely not sufficient to rescue the motility index (**Figure 5M**). Collectively, these results showed that Kif21b is required to fine tune actin dynamics during specific steps of glia-guided locomotion including nucleokinesis and branching of the leading process.

**Figure 5.**
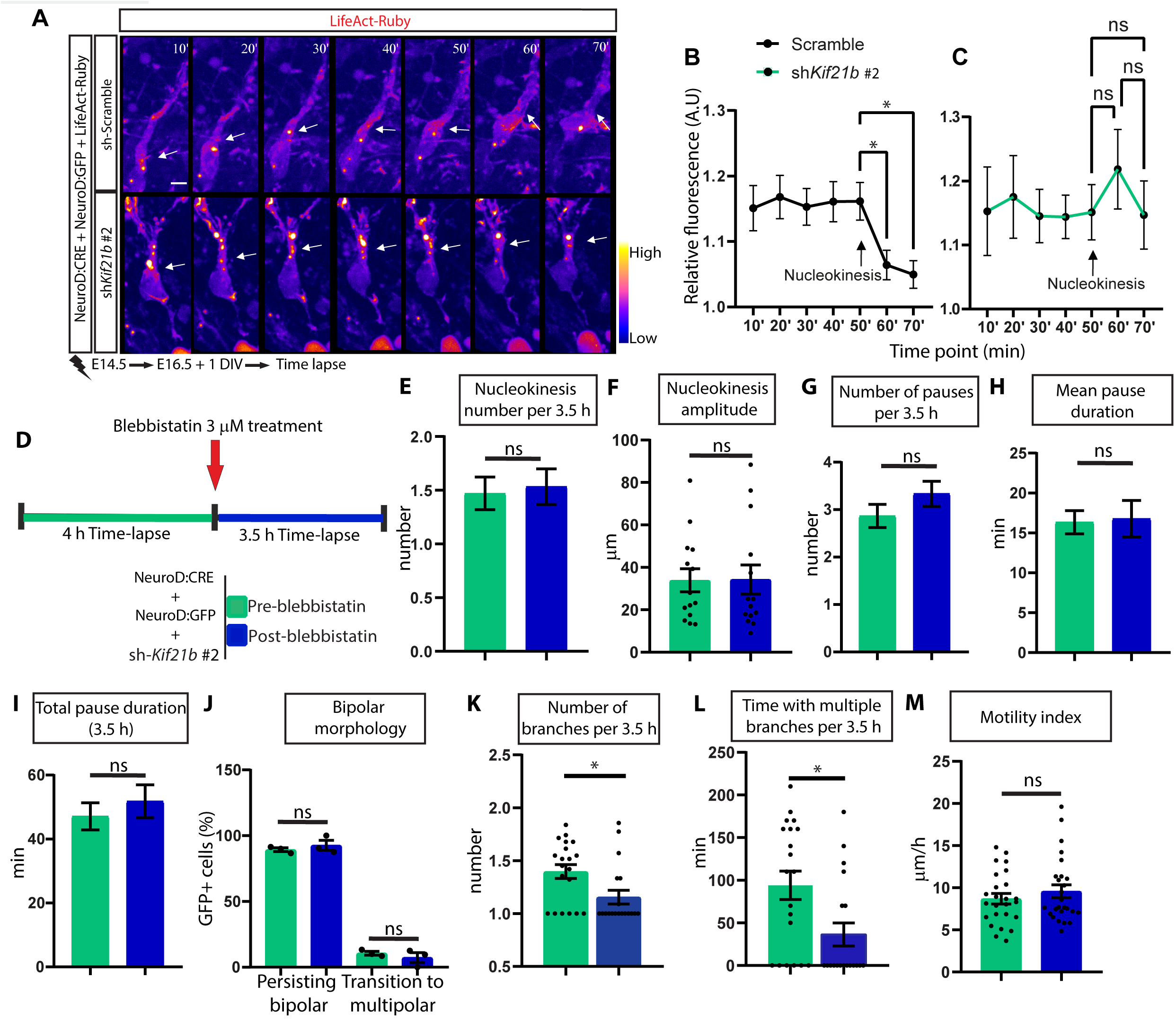
*Kif21b* depletion leads to aberrant actin dynamics in migrating neurons. (**A**) Representative sequence of migrating pyramidal neurons recorded during 5 hours in E16.5 organotypic slices electroporated at E14.5 with NeuroD:Cre, NeuroD:IRES-GFP and pCAGGS-LifeAct-Ruby together with Cre inducible sh RNA *Kif21b #2* or sh-scramble and cultured for one day. Intensity of the LifeAct-Ruby is shown by ImageJ fire look up table. Scale bar: 10 μm. Arrows points towards the proximal part of the leading process where actin accumulation severely dropped after nucleokinesis in control condition. (**B, C**) Quantification of the relative LifeAct-Ruby fluorescence (A.U.) at the proximal part of the leading process in the LifeAct-Ruby+ control (**B**) or *Kif21b* depleted neurons (**C**) before and after nucleokinesis (pointed by an arrow). Data were analysis by fitting a mixed-model, Tukey’s multiple comparisons test. ns, non-significant; *P < 0.05. Number of cells analysed: sh-scramble, n= 22 and sh-*Kif21b #2*, n=24. (**D**) Schematic representation of the rescue experimental approach with Blebbistatin. E16.5 organotypic slices electroporated at E14.5 by NeuroD:Cre, NeuroD:IRES-GFP and the Cre inducible shRNA *Kif21b #2* were imaged by time-lapse microscopy for 4 hours (h), followed by Blebbistatin incubation, and subsequent imaging for an additional 3.5 hours time-lapse. (**E-J**) Analysis (means ± s.e.m.) of the frequency (**E**) and the amplitude (μm) (**F**) of nucleokinesis as well as the average number of pauses per 3.5h (**G**), the mean pause duration (min) (**H**), the total pausing time (min) (**I**) and the percentage of GFP+ neurons converting to multipolar stage (**J**) pre and post-Blebbistatin treatment indicating that Kif21b-induced effect on nucleokinesis and pausing are independent of non-muscle myosin 2 activity. (**K-L**) Quantification (means ± s.e.m.) of branching parameters before and after Blebbistatin treatment, show rescue of number of branches (**K**) and time spent with multiple branches (min) (**L**) upon non-muscle myosin 2 inhibition. (**M**) Quantification (mean ± s.e.m.) of motility index (μm**/**h). Significance was calculated by unpaired t-test. ns, non-significant; *P < 0.05. Number of cells analyzed from at least 3 embryos: (**E,F**): pre-blebbistatin, n=17 and post-blebbistatin, n=15. (**G-I**) pre-blebbistatin, n=45 and post-blebbistatin, n=33; (**K-L**) pre-blebbistatin, n=20 and post-blebbistatin, n=19; (**M**) pre-blebbistatin, n=25 and post-blebbistatin, n=26. Numbers of embryos analyzed: (**J**) pre-blebbistatin and post-blebbistatin, n=3 embryos.

## Discussion

Our findings uncover physiological roles for Kif21b during radial migration of projection neurons. Although it is dispensable for the new-born pyramidal neurons to reach the intermediate zone and convert to a bipolar shape, Kif21b paces the glia-guided locomotion by controlling nucleokinesis, dynamic branching and pausing of migrating neurons. To date, several members of the kinesin superfamily have been shown to play critical functions during radial migration (Akkaya et al., 2021; Broix et al., 2016; Carabalona et al., 2016; Falnikar et al., 2011; Falnikar et al., 2013; Gilet et al., 2020; Li et al., 2022; Muralidharan et al., 2022; Qian et al., 2022; Sapir et al., 2013; Yu et al., 2020). Yet, unlike Kif21b, many of them regulate the initial multipolar-to-bipolar transition (Broix et al., 2016; Falnikar et al., 2013; Gilet et al., 2020; Qian et al., 2022; Sapir et al., 2013; Yu et al., 2020). Noteworthy, dynamic analysis of each stage of migration (multipolar-to-bipolar transition or bipolar locomotion) has not been performed systematically, preventing the exact delineation of the functions of those kinesins during radial migration.

Our study unravels the molecular mechanisms underlying Kif21b functions in migrating neurons. Several lines of evidence suggest that Kif21b regulates different phases of migration through distinct functional domains. First, our complementation experiments revealed that: i) although the N-terminal motor domain is partially required for Kif21b to promote neuronal migration, its ability to hydrolyze or bind ATP is nonessential; and ii) the C-terminal WD40 repeat (WDR) domain that likely binds cargoes (Marszalek et al., 1999) is not necessary to promote migration. This demonstrates that the N-terminal domain, independently of its motility, and the coiled-coil central region are both necessary but, individually, not sufficient for Kif21b functions in pyramidal neurons during migration. Second, rescue analysis with the truncated Kif21bΔMD protein showed that the N-terminal domain is dispensable for the processivity of locomoting neurons but necessary to maintain the bipolar morphology and to limit pausing during locomotion. It suggests that the N-terminal domain and the coiled-coil domain ensure the functions of Kif21b in neuronal polarization and active locomotion, respectively. As the N-terminal domain is essential for the inhibition of MT growth (Masucci et al., 2022; Taguchi et al., 2022), it is tempting to postulate that Kif21b maintains bipolar morphology through the regulation of MT polymerization. Indeed, neuronal polarization largely relies on remodeling of the MTs cytoskeleton (Lasser et al., 2018; Sakakibara et al., 2013) and stabilization of MTs instructs neuronal polarization (Schelski and Bradke, 2022; Witte et al., 2008). Overgrowth of MTs induced by loss of Kif21b (Hooikaas et al., 2020; Masucci et al., 2022; Muhia et al., 2016) could therefore compromise the bipolarity of locomoting neurons. Such hypothesis should be further tested by examining the ability of MT-depolymerizing drugs, like vinblastine, to rescue bipolar morphology of *Kif21b*-depleted migrating neurons (Hooikaas et al., 2020). In line with a MT-dependent effect, we showed that Blebbistatin treatment does not rescue the faulty bipolar-to-multipolar conversion observed upon *Kif21b* knockdown, suggesting that the role of Kif21b in maintaining bipolar morphology is independent of actomyosin contraction. However, as the N-terminal region of Kif21b binds to the unconventional Myosin Va in a neuronal activity dependent manner (Gromova et al, companion manuscript), one cannot exclude that Kif21b cooperates with Myosin Va to regulate actin dynamics in bipolar neurons, although, to date, neuronal activity has been shown to facilitate the initial step of projection neurons migration -the multipolar-to-bipolar conversion - unlikely regulated by Kif21b (**Figure 2**) (Ohtaka-Maruyama et al., 2018). The central coiled coil domain contains secondary MT-binding domain that promotes MT assembly and limits catastrophe frequency, although it is unclear whether it could act alone or exclusively in combination with the other secondary MT-binding domain within the WD40 tail (Ghiretti et al., 2016; Masucci et al., 2022; van Riel et al., 2017). Further work is required to delineate the function of each MT-binding domains in migrating neurons. Third, our *in vitro* co-sedimentation assays of pre-polymerized actin with purified recombinant Kif21b revealed short fragments of Kif21b that might specifically bind to actin (data not shown). Interestingly, those Kif21b fragments encompass all the functional domains of Kif21b, suggesting that Kif21b binds actin through multiple domains, including its central region. Accordingly, our attempts to precisely map Kif21b actin-binding domain(s) using immunoprecipitation of various Kif21b truncated proteins that either lack or only encompass the motor domain, regulatory coiled-coil domain (rCC), coil-coiled domain 1 or 2 or WD40 region were inconsistent and suggest that Kif21b binding to actin filaments requires specific structural conformation involving distinct and potentially distant protein domains. To date, our knowledge about the precise domains of binding to MT, actin or Myosins is limited, with only large domains identified (Ghiretti et al., 2016; Masucci et al., 2022; van Riel et al., 2017) (Gromova et al, companion manuscript). In addition, the high overlap between those domains raises the possibility of a competitive binding of Kif21b to actin and microtubule networks. Refinement of the domains through structural analysis of Kif21b either bound to actin or microtubule would help, in the future, to understand the cooperative or competitive function(s) of Kif21b on both actin and microtubule cytoskeletons.

Our data provide the first evidence for a direct role of a kinesin on actin cytoskeleton in mammals. Indeed, to date, only a plant- and a *Dictyostelium*-specific kinesin (Iwai et al., 2004; Preuss et al., 2004) were shown to bind actin. Here, we first ascertained a direct binding of Kif21b to actin filaments by super-resolution imaging of cortical neurons in culture and by co-sedimentation assay with purified recombinant proteins (**Figure 4**). This validates previous proteomic data suggesting binding of actin and actin binding partners to exogenous Kif21b purified from HEK293T cells (van Riel et al., 2017). Then, investigations of the role of Kif21b on actin dynamics revealed that Kif21b favors the initial step of actin polymerization, the nucleation, but not the subsequent elongation of actin filaments, suggesting that Kif21b facilitates the formation of actin seeds but does not regulate, at least through direct interaction, the elongation of actin filament once formed. Very interestingly, effect of Kif21b on actin assembly highly depends on the stoichiometric ratio between actin and Kif21b molecules (**Figures 4 and S4**). This is reminiscent of the effect of Kif21b on the dynamics of the MT cytoskeleton in cells, with high and low kinesin concentrations leading to opposite effects on MT growth and stability (Ghiretti et al., 2016; Masucci et al., 2022; van Riel et al., 2017). The complex actin network is further built via Arp2/3-mediated nucleation of new actin branches on the side of pre-existing filaments, hence generating dense fast growing entangled actin networks (Firat-Karalar and Welch, 2011). *In vitro* analysis of filament branching using recombinant Arp2/3 complex and its activator pWA does not reveal changes in branch formation in absence or presence of Kif21b (data not shown). This suggests that Kif21b has likely no direct effect on Arp2/3 dependent nucleation in an assay where no competition with actin binding is present. However, Kif21b might modulate actin branching indirectly through its interaction with Arp2/3 (van Riel et al., 2017) or coronin (Figure S4A), that is known to directly regulate Arp2/3 complexes (Gandhi and Goode, 2008). Notably, in a complementary study, Gromova et al showed that Kif21b promotes polymerization and co-localizes with Arp2/3 complexes at actin foci in Cos-7 cells. Altogether, these findings indicate that Kif21b might regulate all steps of actin network dynamics through either direct (nucleation) or indirect (elongation and branching) mechanisms. At the cellular level, except for the Kinesin 6 that has been shown to perturb actin localization during the initial multipolar-to-bipolar transition through unknown mechanisms (Falnikar et al., 2013), very few is known about kinesin and actin interplay in migrating cortical projection neurons. Here we showed, in control neurons, that filamentous actin accumulates ahead of the nucleus before nucleokinesis, in line with previous observation in both cortical projection neurons (Martinez-Garay et al., 2016) and in the cerebellar granule neurons that also undergo glia-guided locomotion (Solecki et al., 2009).

Reminiscent of what has been shown in cerebellar neuron, in which a retrograde actin flow away from the proximal region of the leading process drives nucleokinesis (Solecki et al., 2009), actin concentration at the front of the nucleus drastically drops in cortical projection neurons after nuclear movement. Interestingly, this dynamic actin turnover does not occur in migrating cortical neurons lacking Kif21b, suggesting that Kif21b controls nucleokinesis by controlling actin remodeling. Because non-muscle myosin II (NM2) activity ahead of the nucleus is critical for nucleokinesis (Solecki et al., 2009; Tsai et al., 2007), we hypothesized that Kif21b, through its binding to both actin and NM2, might regulate actomyosin contraction during nuclear forward movement. However, our rescue experiments using Blebbistatin indicate that the faulty nucleokinesis induced by the depletion of *Kif21b* is independent of the perturbation of NM2 activity (**Figure 5**). This further suggests that the high concentration of F-actin observed, after nucleokinesis, in the proximal leading process upon *Kif21b* depletion is rather caused by an excessive stabilization of F-actin than a defect in the retrograde actin flow, that is thought to be NM2 dependent (Solecki et al., 2009). This would imply that Kif21b has dual function on actin assembly and disassembly, possibly mediated through distinct binding domains, reminiscent of the double activities of Kif21b on MT nucleation and stability (Ghiretti et al., 2016; Masucci et al., 2022; van Riel et al., 2017). Rescue experiment with Latrinculin A, that depolymerizes actin filaments (Fujiwara et al., 2018), would help better delineating actin-dependent function(s) of Kif21b during nucleokinesis. Finally, in contrast to nucleokinesis and pausing, we showed that Kif21b likely regulates the dynamic branching of the leading process through regulation of actomyosin contraction. Collectively, these results indicate that Kif21b promote glia-guided locomotion of projection neurons by regulating both nucleokinesis and dynamic branching though distinct non-canonical functions on actin cytoskeleton.

In conclusion, our results indicate that Kif21b plays pleiotropic functions during all steps of the glia-guided locomotion of cortical projection neurons. These cellular functions, driven by distinct protein domains, are independent of the canonical processivity of Kif21b and likely rely on regulation of both microtubules and actin cytoskeletons dynamics. As it has been suggested for the plant-specific kinesin GhKCH1 (Preuss et al., 2004) and the *Dictyostelium-*specific kinesin DdKin5 (Iwai et al., 2004), our work opens the question of the roles of Kif21b in the interplay between microtubules and actin during neuronal migration, adding potential novel atypical function of kinesins in the coordination of actin and MT networks during mammalian brain development.

## Methods

### Cloning and plasmid constructs

shRNAs to deplete the expression of *Kif21b* were directed against the coding sequence 3390-3410 (NM_001252100.1) (sh-*Kif21b #1*) or the 3’-UTR (sh-*Kif21b #2*) and cloned in the pCALSL-mir30(Matsuda and Cepko, 2007) backbone vector as described in (Asselin et al., 2020). Various truncated Kif21b constructs were generated by Sequence and Ligation Independent Cloning (SLIC) as follows: Kif21bΔMD (Δ9-371, motor domain), Kif21bΔATP (Δ87-94, ATP binding site), Kif21bΔWD40 (Δ1308-1639, WD40 domain). The c.288 C>T substitution that abolishes Kif21b mobility was created from WT CDS by site-directed mutagenesis to generate the Kif21b-T96N construct. Wild-type mouse *Kif21a* CDS (NM_001109040.2) was isolated from E18.5 cDNA mouse cortices by PCR and subcloned by restriction-ligation into the NeuroD:IRES-GFP plasmid (Hand and Polleux, 2011). The plasmid pCAGGs-PACT-mKO1 bearing the pericentrin-AKAp450 centrosomal targeting (PACT) domain fused to Kusabira Orange was kindly (Konno et al., 2008) was used to label centrosome and pCAGGS-LifeAct-Ruby (Riedl et al., 2008) was used for F-actin labeling. A Cre/loxP conditional cherry expressing plasmid pCAG-loxPSTOPloxP-Cherry was used for time-lapse imaging of migrating interneurons. Plasmid DNAs used in this study were prepared using the EndoFree plasmid purification kit (Macherey Nagel).

### Mice

All animal studies were conducted in accordance with French regulations (EU Directive 86/609 – French Act Rural Code R 214-87 to 126) and all procedures were approved by the local ethics committee and the Research Ministry (APAFIS#15691-201806271458609). Mice were bred at the IGBMC animal facility under controlled light/dark cycles, stable temperature (19°C) and humidity (50%) condition and were provided with food and water *ad libitum*.

Timed-pregnant wild-type (WT) CD1 (Charles River Laboratories) females were used for *In Utero* electroporation (*IUE*) at embryonic day 14.5 (E14.5). NMRI (WT) mice (Janvier Labs) were used for growth cone extraction and actin preparation assays. *Kif21b*^tm1a(KOMP)Wtsi^ were obtained from UC Davis/ KOMP repository. *Kif21b*^Flox/Flox^ conditional KO mice were generated using the International Mouse Phenotyping Consortium targeting mutation strategy (Skarnes et al., 2011) by crossing *Kif21b*^tm1a(KOMP)Wtsi^ with a Flipase recombinase mouse line (FlpO-2A-eYFP) (Birling et al., 2012). Genotyping was done as previously described in (Asselin et al., 2020).

### *In Utero* Electroporation

Timed-pregnant mice were anesthetized with isoflurane (2 L per minute (min) of oxygen, 4% isoflurane in the induction phase and 2% isoflurane during surgery: Tem Sega). The uterine horns were exposed, and a lateral ventricle of each embryo was injected using pulled glass capillaries (Harvard apparatus, 1.0OD*0.58ID*100mmL) with Fast Green (1 µg/µl; Sigma) combined with different amounts of DNA constructs using a micro injector (Eppendorf Femto Jet).

Embryos were injected with 1 µg/µl of NeuroD:Cre-IRES-GFP vector and 0.5 µg/µl NeuroD:IRES-GFP vector (empty, WT or mutated for Kif21b or WT-Kif21a construct) together with 3 µg/µl of either Cre inducible pCALSL-miR30-shRNA-*Kif21b* or pCALSL-miR30-sh-scramble plasmids. For rescue experiments and time-lapse imaging of actin dynamics, 1 µg/µl of truncated Kif21b constructs (or empty plasmid as control) or 3 µg/µl of pCAGGS-LifeAct-Ruby plasmid (Riedl et al., 2008) were co-injected with the mentioned plasmids. Plasmids were further electroporated into the neuronal progenitors adjacent to the ventricle by discharging five electric pulses (40V) for 50 ms at 950 ms intervals using electrodes (diameter 3 mm; Sonidel CUY650P3) and ECM-830 BTX square wave electroporator (VWR international). After electroporation, embryos were placed back in the abdominal cavity and the abdomen was sutured using surgical needle and thread. Pregnant mice were sacrificed by cervical dislocation, and embryos were collected at E16.5 or E18.5.

### *In vivo* migration analysis

#### Mouse brain collection, immunolabeling

E16.5 and E18.5 embryos were sacrificed by head sectioning and brains were fixed in 4% paraformaldehyde (PFA, Electron Microscopy Sciences) in Phosphate buffered saline (PBS, HyClone) overnight at 4°C. Dissected brains were embedded in 4% low melt agarose (Bio-Rad), and sectioned with vibratome (Leica VT1000S, Leica Microsystems) in 60 µm slices. Free floating sections were maintained in cryoprotective solution (30 % Ethyleneglycol, 20% Glycerol, 30%DH2O, 20% PO_4_ Buffer) at -20°C until the time of use. Alternatively, for Cux1 immunostaining, after fixation, brains were rinsed and equilibrated in 20% sucrose in PBS overnight at 4°C, embedded in Tissue-Tek O.C.T. (Sakura), frozen on dry ice and coronal sections were cut at the cryostat (18 µm thickness, Leica CM3050S) and maintained at -80°C until the day of immunolabelling.

Free floating sections were permeabilized and blocked with blocking solution containing 5% Normal Donkey Serum (NDS, Dominic Dutsher), 0.1% Triton-X-100 in PBS. Brain slices were incubated with primary antibodies diluted in blocking solution overnight at 4°C and then rinsed and incubated with secondary antibody diluted in PBS-0.1% Triton for one hour at room temperature (RT). Cell nuclei were stained using DAPI (1mg/ml Sigma). Slices were mounted in Aquapolymount mounting medium (Polysciences Inc). For cryosections immunolabeling, antigen retrieval was performed by incubating the sections in boiling citrate buffer (0.01 M, pH 6) for 10 minutes, followed by standard immunostaining procedure. All primary and secondary antibodies used for immunolabeling are listed in **Table S3**.

#### Images acquisition and analysis

Cell counting were done in at least three different brain slices from of at least three different embryos for each condition. After histological examination, only brains with comparative electroporated regions and efficiencies were conserved for quantification.

Images were acquired in 1024x1024 mode using confocal microscope (TCS SP5; Leica) at 20x magnification (z stack of 1,55 μm) and analyzed using ImageJ software. Cortical wall areas (UpCP: upper cortical plate, LoCP: lower cortical plate, IZ/SVZ: intermediate zone, subventricular zone/ventricular zone) were identified according to cell density (nuclei staining with DAPI). The total number of GFP-positive cells in the embryonic brain sections was quantified by counting positive cells within a box of fixed size and the percentage of positive cells in each cortical area was calculated.

### Organotypic slices culture and real-time imaging

#### Organotypic slices culture

*IUE* was performed in CD1 mice as indicated above, E16.5 mouse embryos were killed by head sectioning, and brains were embedded in 4% low-melt agarose (BioRad) diluted in HBSS (Hank’s Balanced Salt Solution, ThermoFisher Scientific) and sliced (300 μm) with a vibratome (Leica VT1000S, Leica Microsystems). Brain slices were cultured 16 to 24 hours in semi-dry conditions (Millicell inserts, Merck Millipore), in a humidified incubator at 37 °C in a 5% CO_2_ atmosphere in wells containing Neurobasal medium supplemented with 2% B27 (Thermofisher #17504044), 1% N2 (Thermofisher #17502048), and 1% penicillin/streptomycin (Gibco, Life Technologies). Slice cultures were placed in a humidified and thermo-regulated chamber maintained at 37 °C on the stage of an inverted confocal microscope.

#### Time lapse recordings

Time-lapse confocal microscopy was performed with a Leica SP8 scanning confocal microscope equipped with a 25X water immersion objective / N.A. 0.95 and a humified incubation chamber (37°C, 5% CO_2_). 25 successive ‘z’ optical plans spanning 50 μm were acquired every 30 minutes for 10 hours. Time-lapse acquisitions of actin dynamics in migrating neurons were performed in a confocal Spinning Disk CSU-W1 microscope using a 25X / N.A. 0.95 water immersion objective equipped with a humified incubation chamber, with acquisitions at 10 minutes intervals for 5 hours. Rescue analysis of neuronal migration with Blebbistatin were done using the same experimental setup except that slices were recorded for 4 hours with 10 minutes intervals, then treated with 3 µM Blebbistatin (abcam, #ab120425) by addition of the drug to the culture media and then recorded for another 3.5 hours.

#### Analysis of time lapse sequences

All sequences were analyzed using ImageJ by adjusting time-interval in the plugin “Manual tracking”, as well adjusting the pixel width with the x/y calibration corresponding to the analyzed sequence. Migrating neurons were manually tracked individually during the time-lapse and the data collected were analyzed for migration velocity calculating the mean velocity during the entire time-lapse acquisition; number of pauses was calculated by counting the number of times that the neuron stopped moving (velocity = 0); mean and total duration of pauses were calculated by the mean and total number of minutes in which the neuron spent without moving (velocity=0), respectively; and motility index was calculated as the mean velocity during the entire time-lapse acquisition subtracting the pausing time (velocity=0). Analysis of the proportions of multipolar-bipolar, persisting multipolar, bipolar-multipolar and persisting bipolar neurons were done by the identification of migrating neurons with either multipolar or bipolar morphology and the follow-up of morphological changes (or absence of them) during the entire duration of the time-lapse. Swelling and nucleokinesis were counted for each neuron during the time-lapse, as well as the time each neuron took to complete a nucleokinesis. Additionally, nucleokinesis amplitude was measured by tracking the maximum length of the movement of the soma of each bipolar neuron. Branching in the leading process was quantified by counting the number of branches of each neuron in every time point of the time-lapse, followed by the calculation of the mean number of branches and the sum of the minutes that the neuron spent with 2 or more branches. Length of the branches was calculated considering the time-point in which the neuron extended at its maximum length the leading process, or main leading process for branching neurons, in which case, the length of all the branches was counted for that time-point.

LifeAct fluorescence was determined by delimitating the area of the proximal part of the leading process where the LifeAct signal showed increasing levels and measuring the mean grey values of the delimited region during for each time-point of the time-lapse. For each measure, normalization was done with the mean grey fluorescence with a ROI in the soma, where values of fluorescence were constant. Time-lapse analyses were done in at least three different brain slices from different embryos in at least three independent experiments for each condition.

### *Ex-vivo* electroporation of medial ganglionic eminence (MGE) explants and time lapse imaging of migrating interneurons

*Ex-vivo* electroporation of MGE and time-lapse imaging has been done as previously described (Tielens et al., 2016). The heads of E13.5 Dlx5,6 Cre-GFP (Stenman et al., 2003) embryos were harvested and the cortex removed in order to expose the ganglionic eminences. Plasmids to overexpress Cre inducible pCALSL-miR30-shRNA-*Kif21b* or pCALSL-miR30-sh-scramble were used at a concentration of 3 µg/µl and were directly injected in the MGEs. These plasmids were co-electroporated with a Cre/loxP conditional cherry expressing plasmid used at a concentration of 1 µg/µl. Electroporation conditions: 50V, 5 pulses of 50ms duration spaced by 1s interval. Electroporated MGEs were then cultured on cortical cell feeder seeded on glass-bottom petri-dish (MatTek, Ashland, USA). Co-cultures were placed in a humidified and thermo-regulated chamber maintained at 37°C on the stage of an inverted confocal microscope. Time-lapse confocal microscopy was performed with a Leica SP8 scanning confocal microscope with a 25X objective. Images of living cherry-expressing migrating interneurons were acquired every 5 minutes for 4 hours. Migration velocity was analyzed adjusting time interval in the plugin “Manual tracking”, as well as the x/y calibration with the pixel width of the analyzed sequence. Migrating neurons were manually tracked individually during the time-lapse and the data collected was analyzed for migration velocity. Analyses were done in at least three different embryos in two independent experiments for each condition.

### Neuronal culture, fixation and immunostaining

CD1 mice cortices from E15.5 mice were dissected and collected in cold PBS supplemented with BSA (3 mg/mL), MgSO4 (1 mM, Sigma), and D-glucose (30 mM, Sigma). Cortices were dissociated in Neurobasal media containing papain (20U/mL, Worthington) and DNase I (100 μg/mL, Sigma) for 20 minutes at 37°C, washed 5 minutes with Neurobasal media containing Ovomucoid (15 mg/mL, Worthington), and manually triturated in OptiMeM supplemented with D-Glucose (20mM). 220 000 cells were plated in Neurobasal Supplemented media supplemented with B27 (Thermofisher #17504044), L-glutamine (2 mM) (Thermofisher #25030-123), and penicillin (5 units/mL) – streptomycin (50 mg/ml) (Thermofisher #15140-130) on round coverslips of N1.H5, 18 mm diameter, pre-coated with Poly-L-Lysine (1 mg/ml). Neurons were incubated in an incubator with controlled CO_2_ (5%) and temperature (37°C).

At *Day In Vitro* 2 (DIV2) neurons were incubated with pre-warmed extraction buffer containing 0.25% triton, 0.1% glutaraldehyde, in PEM buffer (80 mM PIPES, 5 mM EGTA, 2 mM MgCl_2_, pH 6.8) for 30 seconds and fixed in a pre-warmed fixing solution containing 0.25% Triton, 0.5% Glutaraldehyde, in PEM buffer. Glutaraldehyde was quenched with 0.1 % NaBH_4_ prepared with phosphate buffer (PB) for 7 minutes and cells were washed twice with PB. Blocking was performed by incubation with 0.22% gelatine for 2 hours at RT with gentle shaking, primary antibody diluted in blocking buffer was incubated at 4°C overnight. Secondary antibody diluted in blocking buffer was incubated one hour and washed three times with PB. To label F-actin, phalloidin coupled to Alexa Fluor Plus 647 (Invitrogen, A30107) was incubated overnight for aquapolymount mounting for confocal imaging or incubated until the day of acquisition for super resolution microscopy.

Neurons in Figure 4F were imaged using the LiveSR mode of Confocal Spinning Disk CSU-X1 “Nikon” equipped with a Photometrics Prime 75B camera and an APO TIRF 100x/ N.A. 1.49 oil objective controlled by Metamo0rph 7.10. software.

### SMLM imaging protocol

#### Setup

SMLM was performed with a home-build setup based on an inverted microscope (Eclipse TiE Nikon) equipped with 100x/1.49 NA oil-immersion objective (Apochromat TIRF, Nikon), Perfect Focus and driven with µmanager software (Edelstein et al., 2014). The microscope was equipped with a laser line 532 and 642 nm lasers (Oxxius). Excitation was done in Total Internal Reflection Fluorescence (TIRF) mode to excite only the sample near the surface (< 200 nm). A multiband dichroic mirror was used to filter emission from the sample (FF560/659-Di01, Semrock) and notch filters were used to remove scattered laser light (532 nm and 642 nm StopLine single-notch filters: NF01-532U and NF03-642, Semrock). Spectral discrimination of the two different fluorescent probes was achieved by splitting the fluorescence emission on the camera chip by an optical separator (Gemini, Hammatsu associated to a dichroic mirror (FF 640-Di01 Semrock and respectively a LP 532 (Semrock) and a LP 647 (Semrock) for the top and bottom chip areas)). All images were recorded using an EMCCD camera ((C9100-23B ImagEM, Hamamatsu) with 16 x 16 µm pixels), using 240 x 240 top region on the camera for AlexaFluor 555 labelling and 240 x 240 bottom region for AlexaFluor 647 labelling. Chromatic aberration was corrected by imaging TetraSpek fluospheres excited with a low laser power (532nm and 647 nm). A raster scan of a single bead was performed to record a reference image and determined, for each frame, its localization with the help of DOM ImageJ plugin. This plug-in was also used to determine and corrected the optical and chromatic aberration between the two channels.

#### Acquisition imaging procedure for SMLM

After neurons were cultured, fixed and immunostained as explained above, the coverslips were placed in an observation chamber (Ludin chamber, LIS) and recovered by fresh 700 μL of imaging solution (prepared just before the use by mixing 7 μL of GLOX buffer (56 mg/mL Glucose oxidase, 3.4 mg/mL catalase, 8 mM Tris 10, 40 mM NaCl 50), 70 μL MEA buffer (1 M MEA in 0.25 N HCl) and 620 μL of buffer containing 50 mM Tris (pH 8.0), 10 mM NaCl and 10% Glucose) to control the redox environment. Acquisition was performed by adjusting the laser power to 50 mW at the objective for both lasers that results in 2 kW/cm2 excitation intensity and acquisition time to get single fluorophores bursts with the optimal signal-noise ratio. For the dual-color image acquisition, a sequential approach was used with a first dSTORM recording with Alexa Fluor 647 labelling followed by a second dSTORM recording with the Alexa Fluor 555 labelling into two individual image stacks (more than 30 000 frames per stack).

#### SMLM Data analysis

The two sets of images were read by an imageJ macro, which extracts the appropriate ROI and corrects the chromatic aberrations. Single-molecule localizations in both extract datasets were calculated using the thunderSTORM algorithm (Ovesny et al., 2014). The following parameters were used to find and fit the signal of each particle: image filtering – Difference-of-Gaussians filter (sigma 1 = 1.0 and sigma 2 = 1.6); approximated localization of molecules – Local maximum (peak intensity threshold: std (Wave.F1), connectivity: 8-neighbourhood); sub-pixel localization of molecules – Integrated Gaussian (fitting radius: 4 px, fitting method: Least squares, initial sigma:1.3 px). The reconstructed images of both channels were combined to generate a two colour-image. To calculate the distances between actin and Kif21b clusters, the localization data of the two channels (obtained with ThunderStorm) were analyzed with the PoCA software (https://github.com/flevet/PoCA, plug-in developed by Floriant Levet). Single molecule localization coordinates were used to compute a Voronoï tessellation, in order to partition the image space in polygons of various sizes centered on each localized molecule (Levet et al., 2015). Using this space-partitioning framework, first-rank densities δ_i_^1^ of the molecules were computed. Segmentations were performed thresholding these density features with a threshold of 5 times (resp. 2 times) the average density δ of the whole dataset for the Kif21b clusters (resp. actin filaments). Finally, we defined the shortest distance between the Kif21b clusters and the actin filaments as the distance between the clusters’ centroid and the closest point of the actin filament borders.

### Cell culture and transfections

All cells used in this study are provided by the cell culture platform of the IGBMC (Illkirch), are guaranteed mycoplasma free (PCR test Venorgem) and have not been authenticated. Mouse neuroblastoma N2A cells were cultured in DMEM (1g/L glucose) (Dulbecco’s Modified Eagle Medium, GIBCO) supplemented with 5% Fetal Calf Serum (FCS) and Gentamycin 40µg/mL and Human embryonic kidney (HEK) 293T cells were cultured in DMEM (1g/L glucose) (GIBCO) supplemented with 10% Fetal Calf Serum (FCS), penicillin 100 UI/mL, streptomycin 100 µg/mL in a cell culture incubator (5% CO_2_ at 37°C). Both cell lines were transfected using Lipofectamine 2000 (Invitrogen) according to the manufacturer’s protocol. N2A cells were transfected using the different NeuroD:Kif21b-IRES-GFP constructs. HEK293K cells were transfected using sh-scrambled or ShRNA-*Kif21b* #2 together with pEGFP-C1-3’UTR KIF21B or sh-scrambled or ShRNA-*Kif21b* #1 together with pEGFP-C1-WT-Kif21b. Expression of transfected genes was analyzed 48 hours after transfection by RT-qPCR or immunoblotting.

### Protein extraction and western blot

Proteins from mouse cortex (E18.5) or transfected cells (N2A or HEK 293T) were extracted as follows: cells were lysed in RIPA buffer (50 mM Tris pH 8.0, 150 mM NaCl, 5 mM EDTA pH 8.0,1% Triton X-100, 0.5% sodium deoxycholate, 0.1% SDS) supplemented with EDTA-free protease inhibitors (cOmplete™, Roche) for 30 minutes, then cells debris were removed by high speed centrifugation at 4°C for 25 minutes. Protein concentration was measured by spectrophotometry using Bio-Rad Bradford protein assay reagent. Samples were denatured at 95°C for 10 minutes in Laemmli buffer (Bio-Rad, #1610747) with 2% β-mercaptoethanol (Sigma, #M3148) and then resolved by SDS–PAGE and transferred onto PVDF membranes (Merck, #IPVH00010). Membranes were blocked in 3% milk in PBS buffer with 0.1% Tween (PBS-T) and incubated overnight at 4°C with the antibodies indicated in **Table S3** diluted in blocking solution. Membranes were washed 3 times in PBS-T and incubated at room temperature for 1 hour with the HRP-coupled secondary antibodies indicated in **Table S3**, followed by 3 times PBS-T washes. Visualization was performed by quantitative chemiluminescence using SuperSignal West Pico PLUS Chemiluminescent Substrate (ThermoFisher, #34580). Signal intensity was quantified using ImageJ.

### RNA extraction, cDNA synthesis and RT–qPCR

Total RNA was extracted from the ganglionic eminences of WT NMRI mouse embryos at different time points of development, with TRIzol reagent (Thermo Fisher Scientific). Kif21b ex2-3 (Fwd sequence: AAGGCTGCTTTGAGGGCTAT, Rev sequence: AAAGCCGGTGCCCATAGTA) were used to target mouse *Kif21b* cDNA and actin (Fwd: TATAAAACCCGGCGGCGCA ; Rev: TCATCCATGGCGAACTGGTG) as housekeeping gene normalizer. RT–qPCR was performed in a LightCycler PCR instrument (Roche) using SYBR Green Master Mix (Roche).

### Immunoprecipitation

For immunoprecipitation (IP) experiments, E18.5 brain cortices were lysed using 200 µl of IP buffer (Thermofisher IP Kit (#8788) (1% NP40, 5% Glycerol, 1 mM EDTA, 150 mM NaCl, 25 mM Tris-HCl pH 7.4)) supplemented with EDTA-free protease inhibitors (cOmplete, Roche) and proteins were extracted as explained above. 200 µg of protein were incubated with Kif21b antibody (0.20 µg), actin antibody (2 µg) or corresponding control IgG at 4°C overnight with 5 µL of pre-washed Pierce Protein A/G Magnetic Beads (Thermo-Scientific, # 88802) (see **Table S3** for antibodies references). Magnetic beads were collected with a magnetic stand (Invitrogen, 12321D), washed twice with IP buffer and proteins were eluted with 20 µL of Low pH elution buffer (Thermofisher, #1862619) at room temperature for 10 minutes. Neutralization buffer (Tris pH 8, 1.0 M) and 20 µL of 2x Laemmli Elution Buffer containing 2% β-mercaptoethanol was added to the proteins. Samples were subjected to SDS-PAGE for Western Blot analysis.

### Mass spectrometry

#### Liquid digestion

For mass spectrometry analysis, IP was performed as described above, except that for the protein incubation step, 2 mg of protein,100 μL of pre-washed beads and 3 μg of Kif21b or control IgG antibody were used. After Kif21b elution with 45 µL of low pH elution buffer (Thermofisher, #1862619), the same volume of 2x Laemmli Elution Buffer containing 2% β-mercaptoethanol was added to the sample and the proteins were conserved at -20°C until the analysis. Protein mixtures were precipitated with trichloroacetic acid (TCA) overnight at 4°C, pellets were washed twice with 1 mL cold acetone, dried and dissolved in 8 M urea in 0.1 mM Tris-HCl pH 8.5 for reduction (5 mM TCEP, 30 minutes), and alkylation (10 mM iodoacetamide, 30 minutes). Double digestion (LysC - Trypsin) was performed by incubating the proteins first with endoproteinase Lys-C (Wako, #125-05061,)) in 8 M urea for 4h at 37°C followed by an overnight trypsin digestion (Promega #V511A,) in 2 M urea at 37°C. Peptide mixtures were then desalted on C18 spin-column and dried on Speed-Vacuum before LC-MS/MS analysis.

#### LC-MS/MS Analysis

Samples were analyzed using an Ultimate 3000 nano-RSLC (Thermo Scientific, San Jose California) coupled in line with a LTQ-Orbitrap ELITE mass spectrometer via a nano-electrospray ionization source (Thermo Scientific, San Jose California). One microgram of tryptic peptides (in triplicate) were loaded on a C18 Acclaim PepMap100 trap-column (75 µm ID x 2 cm, 3 µm, 100Å, Thermo Fisher Scientific) for 3.5 minutes at 5 µL/minute with 2% ACN, 0.1% FA in H_2_O and then separated on a C18 Accucore nano-column (75 µm ID x 50 cm, 2.6 µm, 150Å, Thermo Fisher Scientific) with a 90 minutes linear gradient from 5% to 35% buffer B (A: 0.1% FA in H_2_O / B: 99% ACN, 0.1% FA in H_2_O) followed by a regeneration step (90% B) and a equilibration (5%B). The total chromatography was 120 minutes at 200 nL/minute and at 38°C. The mass spectrometer was operated in positive ionization mode, in Data-Dependent Acquisition (DDA) with survey scans from m/z 350-1500 acquired in the Orbitrap at a resolution of 120,000 at m/z 400. The 20 most intense peaks (TOP20) from survey scans were selected for fragmentation in the Linear Ion Trap with an isolation window of 2.0 Da and were fragmented by CID (Collision-Induced Dissociation) with normalized collision energy of 35%. Unassigned and single charged states were rejected. The Ion Target Value for the survey scans (in the Orbitrap) and the MS2 mode (in the Linear Ion Trap) were set to 1E6 and 5E3 respectively and the maximum injection time was set to 100 ms for both scan modes. Dynamic exclusion was used. Exclusion duration was set to 20 s, repeat count was set to 1 and exclusion mass width was ± 10 ppm.

### MS Data Analysis

Proteins were identified with Proteome Discoverer 2.4 software (PD2.4, Thermo Fisher Scientific) and *Mus Musculus* proteome database (Swissprot, reviewed, release 2020_11_20). Precursor and fragment mass tolerances were set at 7 ppm and 0.6 Da respectively, and up to 2 missed cleavages were allowed. Oxidation (M) was set as variable modification, and Carbamidomethylation (C) as fixed modification. Peptides were filtered with a false discovery rate (FDR) at 1%, rank 1. Proteins were quantified with a minimum of 2 unique peptides based on the XIC (sum of the Extracted Ion Chromatogram). The quantification values were exported in Perseus for statistical analysis involving a log[2] transform, imputation, normalization before Volcano plots (Tyanova et al., 2016).

### F-actin sedimentation assay from embryonic cortices

Microdissected E18.5 cortices (3 to 4 cortices per 500 μl) were lysed in a buffer containing 50 mM PIPES pH 6.9, 50 mM NaCl, 5 mM MgCl_2_, 5 mM EGTA, 5% glycerol, 0.1% NP40, 0.1% Triton X-100, 0.1% Tween 20, 0.1% β-mercaptoethanol, supplemented with EDTA-free protease inhibitors (cOmplete™, Roche) for 10 minutes at 37°C. The lysate was first centrifuged at 2,000 g for 5 minutes to remove nuclei then subjected to a high-speed centrifugation (100,000 g, 1 hour, 37°C) to obtain a clear supernatant (G-actin) and a pellet (F-actin). Pellet was dissolved overnight at 4°C in water containing 10 μM cytochalasin D (Sigma-Aldrich #C8273) at the same volume as the collected supernatant and equal amounts of each fraction were subjected to western blot analysis.

### Growth cone extraction from mouse brain

Isolation of growth cone was performed on WT NMRI mouse cortices at post-natal day (P) 2 as previously described (Li et al., 2013). Five P2 mouse brains were homogenized in 5 mL of homogenization buffer (5 mM Tris-HCl, 0.32 M sucrose, 1 mM MgCl_2_) supplemented with EDTA-free protease inhibitors (cOmplete™, Roche), then centrifuged at 1,660 x g for 15 min at 4 °C. Supernatant were loaded on top of a discontinuous 0.75/1.0/2.33 M sucrose density gradient and centrifuge at 242,000 x g for 60 min at 4°C (Beckman SWTi41 swing rotor). The growth cone depleted fraction was collected between 0.75 M and 1.0 M sucrose. The growth cone enriched fraction was collected at the interface between the load and 0.75 M sucrose, re-suspend in the homogenization buffer and centrifuged at 100,000 *x g* for 40 min at 4°C (Beckman SWTi41 swing rotor). The pellet containing the growth cone were re-suspended in 50 μl of the homogenization buffer. The same volume of each fraction was analyzed by western blot. Non-growth cone membranes, which also contain Golgi membranes, were revealed by the enrichment of Golgi matrix protein (GM130 primary antibody), while the growth cone fraction was revealed by the enrichment of Jip1 protein.

### Purification of human KIF21B overexpressed in BHK21 cells

#### MVA expression vector

A vaccinia virus (strain Modified Vaccinia Ankara - MVA - Bio safe level 1) allowing an inducible expression of human KIF21B in mammalian cells (BHK21) were used. Briefly, KIF21B tagged with 6 His-tag at the C-terminal was integrated by homologous recombination at the HA locus of the MVA viral genome.

#### BHK21 overexpression

For protein production, a 1.8 L preculture of BHK21 C13-2P cells (10^6^ cells/ml) grown in Glasgow’s modified Eagle’s medium (GMEM; Thermo Fisher, MA, USA) supplemented with Bacto Tryptose Phosphate (1.5 g/L), 10% foetal calf serum and 50 µM Gentamycin was infected with approximately 0.1 PFU/cell of recombinant virus and incubated at 37°C. After 48 hours, the infected cells were mixed with 6 L of uninfected cells at 10^6^ cells/mL at a 1:10 ratio (v/v). Overexpression was induced by the addition of 1 mM IPTG to cell mixture and cells were incubated at 37°C in 5% CO_2_ at 75% hygrometry for 24 hours. Cells were pelleted at 2000 g for 20 minutes, washed in PBS and pelleted again at 2000 g for 10 minutes. Cell pellet was stored at -20°C until use.

#### KIF21B-6His purification

Pellets from 5L cells were unfrozen and resuspended in lysis buffer (50 mM HEPES pH7.5, 500 mM NaCl, 1 mM MgCl_2_, 10 mM Imidazole, 0.5% IGEPAL® CA-630, 2 mM β-Mercapto-ethanol supplemented with Roche cOmplete inhibitor cocktail tablets) at a ratio of 15 ml of buffer/g of biomass. Lysis was performed by pulse sonication on ice (5 minutes with pulses 2s ON, 2s OFF). Lysate was then clarified by ultracentrifugation for 1 hour at 100.000g at 4°C. After filtration on cellulose filter 5µm, the supernatant was loaded on HisTrap Excel 5mL column (Cytivia). Column was washed with 10 mM imidazole and proteins were eluted with a gradient from 10 to 500 mM of imidazole. Peak fractions at 280 nm were analyzed on SDS-PAGE and fractions containing the complex were pulled together. The sample was then concentrated using an Amicon Ultra 15 ml with a 100-kDa molecular weight cutoff (MWCO) and further purified using a HiLoad Superdex 200 pg 16/60 column (Cytivia) equilibrated with GF buffer (10 mM HEPES pH7.5, 250 mM NaCl, 1 mM MgCl_2_, 2 mM DTT). 2 fractions containing the purest complex (checked by SDS-PAGE) were pooled together to obtain two pooled recombinant sample, concentrated using an Amicon Ultra 15 ml with a 100-kDa molecular weight cutoff (MWCO) to 0.7 mg/ml, flashed frozen in liquid nitrogen and stored at -80°C. The contaminants found in the two pooled samples can be found in **Table S2**.

### *In Vitro* F-Actin and microtubules co-sedimentation assay

Actin was polymerized according to the manufacturer instructions with some modifications to prepare F-actin at 21 µM. Briefly, actin (Cytoskeleton, #AKL99-A) was resuspended in general actin buffer (5 mM Tris-HCl pH 8.0 and 0.2 mM CaCl2), and incubated 30 minutes at 4°C. Actin polymerization buffer (500 mM KCl, 20 mM MgCl_2_, 10 mM ATP) was added and incubated 1 hour at 24°C.

The binding experiments were performed for the following conditions: Kif21b alone and Kif21b+Actin; using 5 µl of recombinant Kif21b (from fraction 1), 29 µl of polymerized actin and adding actin buffer to complete a volume of 50 µl. The reactions were incubated 30 minutes at 24°C and ultracentrifuged 60 minutes at 60 000 rpm in an Optima MAX-E Ultracentrifuge, using the rotor TLA 100.4. Supernatant was collected and resuspended in 10 µl of 4x Laemmli buffer containing 2% β-mercaptoethanol, while the pellet was resuspended adding 30 μl of milliQ water followed by 30 μl of 2% β-mercaptoethanol. Pellet and supernatant fractions were then analysed by Coomassie staining. After running the samples in a SDS-PAGE gel, the gel was warmed for 40 seconds in a microwave in a solution containing 50% ethanol, 10% acetic acid, and incubated on agitation at RT for 10 minutes. It was then incubated overnight in a solution containing 5% ethanol and 7.5% acetic acid with Coomassie blue. The gels were washed in milliQ water to remove background.

### Actin polymerization assays *In Vitro*

#### Coverslips preparation

Glass coverslips were oxidized with oxygen plasma (30s, 35%, Diener Electronic, cat. ZeptoB) and incubated with 5% BSA in HEPES 10 mM at pH 7.4 for 10 minutes RT and washed with HEPES 10 mM.

#### Actin polymerization of cortices extract

Actin assembly was induced by mixing 7,36 μL of cortex extract with the reaction mixture containing 18 μL fluorescent buffer (15 mM imidazole, pH 7.0, 74 mM KCl, 1.5 mM MgCl_2_, 165 mM DTT, 2 mM ATP, 50 mM CaCl_2_, 5 mM glucose, 30 mg/ml catalase, 155 mg/ml glucose oxidase, and 0.75% methylcellulose.), 2.64 μL of G buffer (2mM Tris-HCl, 0.2 mM Na_2_ ATP, 0.2 mM CaCl_2_, 5 mM DTT), 0,1% BSA and 1 μL of phalloidin 488 nm (A12379,1/200, diluted in Hepes 10 mM). 5 μL of the reaction mixture were immediately put between coated-coverslip and slide and sealed with VALAP (1:1:1 vaseline, lanolin and paraffin).

#### Actin polymerization with purified protein

Actin assembly was induced by mixing TicTac buffer (Farina et al., 2016) (10 mM HEPES,16 mM PIPES, 50 mM KCl, 5 mM MgCl2,1 mM EGTA supplemented with - 2.7 mM ATP, 10 mM DTT, 20 μg/mL Catalase, 3 mg.mL Glucose, 100 μg/mL Glucose Oxydase, 0.25% Methycellulose) with 0,1% BSA and 1/60 000 phalloidin (Fisher Scientific A12379). Kif21b (from fraction 2) was added at a final concentration of 1 nM or 10 snM. The reaction mixture was immediately put in a flow chamber (constitute of clean glass slide, a BSA-coated coverslip, and precut adhesive double tape 50 μm thick) and sealed with VALAP (1:1:1 vaseline, lanolin and paraffin).

#### Image acquisition

Images were taken using an inverted Nikon EclipseTi microscope equipped with a ×100 oil objective (HCX plan APO) and a Photometrics Prime 95B (Teledyne Photometrics). The microscope and devices were driven by MetaMorph (Molecular Devices, Downington, PA). Images of actin polymerization of cortices extract were acquired every 1s with the following parameters: 30% of laser power and 200ms of exposure time. Image of actin polymerization with purified proteins were acquired every 5s with the following parameters: 50% of laser power and 100ms of exposure time.

#### Image Analysis

Filaments were segmented using TSOAX (V0.2.0, default parameters except for Gaussian-std: 2 and by adjusting the ridge threshold) (Xu et al., 2019). The resulting file was treated using a home-made script on R studio (Version 1.4.1106). The results were plot using a Prism 9. The total amount of polymerized actin was calculated as the sum of all filaments detected at each time.

Then, the mean amount of polymerized actin was plotted, the linear regression was calculated, and statistical differences were calculated with GraphPad Prism 9. Mean length of all the detected filaments was measured at each time point. The results were analyzed by fitting the exponential curve and statistical differences was calculated with Prism 9. To quantify actin elongation rate, single filaments were manually tracked: filament length was measured during 90s. The results were plotted, and the linear regression was calculated, and statistical differences was calculated with GraphPad Prism 9.

### Statistical analysis

All statistics were calculated using GraphPad Prism 6 or 9 (GraphPad) and are represented as mean ± s.e.m. The number of experiment repetitions and statistical tests are indicated in the figure legends and also reported in Supplementary data 1. Adjustments made for multiple comparisons, confidence intervals and exact P-values for Figs.1B, 1E; 2E-K, M-P, R-W; 3C, E-J; 4H, J-M; 5B-C, E-M; Supplementary 1A, B, D; Supplementary 2A, B, D; Supplementary 3C; Supplementary 4C, D are reported in Supplementary Data 1.Graphs were generated using GraphPad and images were assembled with Adobe Illustrator CS6 (Adobe Systems).

## Acknowledgments

This work was funded by grants from INSERM (ATIP-Avenir program, J.D.G.), the Fyssen foundation (J.D.G.), the French state funds through the Agence Nationale de la Recherche under the project JCJC CREDO ANR-14-CE13-0008-01 (J.D.G.), and the programme Investissements d’Avenir labelled (ANR-10-IDEX-0002-02, ANR-10-LABX-0030-INRT, to J.D.G., A-C.R. and M.R.), INSERM/CNRS and University of Strasbourg. This work of the Interdisciplinary Thematic Institute IMCBio, as part of the ITI 2021-2028 program of the University of Strasbourg, CNRS and INSERM, was supported by IdEx Unistra (ANR-10-IDEX-0002), and by SFRI-STRAT’US project (ANR 20-SFRI-0012) and EUR IMCBio (ANR-17-EURE-0023) under the framework of the French Investments for the Future Program. L.A. and J.R.A. were funded through the IGBMC PhD program (ANR-10-IDEX-0002-02, ANR-10-LABX-0030-INRT) and Fondation pour la recherche médicale. R.B. is funded by an ANR JCJC granted to A.C.R. (ANR-19-CE13-0005-01). P.T. and J.B. are, respectively, research assistant and assistant professor at the University of Strasbourg. C.B. is funded by CERBM-GIE. L.R. and B.M. are research engineers at CNRS. F.L. is a research engineer at INSERM. J.D.G. is an INSERM investigator. Y.M. is professor at the University of Strasbourg and Institut Universitaire de France. A-C.R. is a CRNS investigator. We thank the Imaging Center of IGBMC (ici.igbmc.fr), in particular, Elvire Guiot and Erwan Grandgirard for their assistance in the imaging experiments. We are grateful to the staff of the mouse facilities of the Institut Clinique de la souris (ICS) and Institut de Génétique et de Biologie Moléculaire et Cellulaire (IGBMC), the staff of the proteomics platform and of the molecular biology service (in particular Thierry Lerouge and Paola Rossolillo) for their involvement in the project. The SMLM imaging was supported by the Imaging Center PIQ-QuESt (https://piq.unistra.fr/). We warmly thank Dr Binnaz Yalcin, Dr Laurent Blanchoin and Christophe Guérin for helpful comments, advice, and reagents. We are also grateful to members of J.D.G. laboratory for discussion and technical assistance.

## Author contribution

J.R.A. and L.A. conceived and designed the experiments, performed the experiments, performed statistical analysis, and analyzed the data related to cellular, and functional studies in mice. P.T. performed *in utero* electroporation. P.T and N.S provided technical assistance. R.B. and A-C.R. conceived, designed, and performed *in vitro* actin experiments. C.B., J.B. and M.R. contributed to the production of recombinant Kif21b proteins. B.M. performed mass spectrometry analysis and analyzed data. L.R. performed SMLM acquisition and performed quantitative analysis using a plugin developed by F.L. Y.M. contributed to SMLM data analysis and interpretation. J.D.G. conceived, coordinated, and supervised the study, designed experiments, analyzed data and wrote the manuscript.

