## Supplementary information for "The kinesin Kif21b regulates radial migration of cortical projection neurons through a non-canonical function on actin cytoskeleton"

Rivera Alvarez *et al*

Content:

- **Figure S1:** Depletion of *Kif21b* in projection neurons impairs migration, related to Figure 1.
- **Figure S2:** *Kif21b* depletion induces defects in tangential migration of interneurons, related to Figure 1.
- **Figure S3:** Overexpression of Kif21b-truncated constructs does not alter radial migration, related to Figure 3.
- **Figure S4:** Kif21b interacts with actin and actin binding proteins and regulate actin dynamics, related to Figure 4.

**Figure S1: Depletion of *Kif21b* in projection neurons impairs migration, related to Figure 1.**

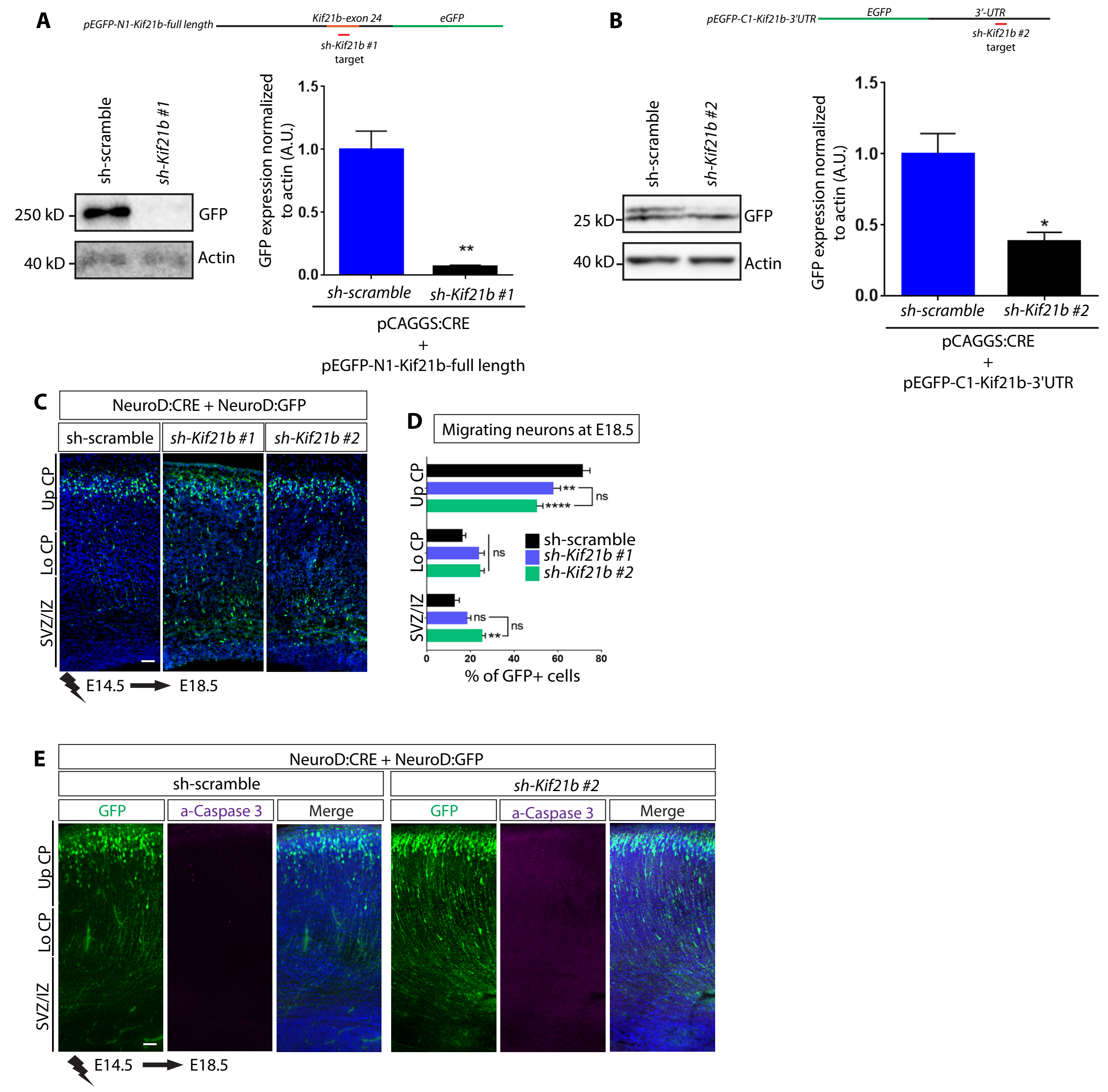


(**A,B**) Western blots of extracts HEK293T cells transfected with the indicated constructs showing the efficiency of the shRNA-Kif21b #1 directed against the coding exon 24 (**A**) and shRNA-Kif21b #2 targetting the 3'UTR (**B**). Data (means ± s.e.m) from 3 independents experiments were analyzed by unpaired two-tailed Student t-test, $^{**}P < 0.005$. (**C**) Coronal sections of E18.5 cortices electroporated with NeuroD:Cre and NeuroD:IRES-GFP together with Cre inducible shRNA *Kif21b* #1, *Kif21b* #2 or sh-scramble showing similar migration defects using two different shRNA sequences. **D** Quantification (means ± s.e.m.) of the distribution of GFP-positive neurons in different regions (Up CP, Upper cortical plate; Lo CP, Lower cortical plate; SVZ/ IZ, subventricular zone / intermediate zone) in all the indicated conditions. Significance was calculated by two-way ANOVA, Bonferroni's multiple comparisons test. ns, non-significant; **P < 0.005; ****P < 0.0001. Number of embryos analyzed: sh-scramble, n=7; sh-*Kif21b* #1, n=8; sh-*Kif21b* #2, n=9. (**E**) Immunolabelling of activated caspase 3 (a-caspase 3, in purple) in E18.5 mouse cortices electroporated at E14.5 with NeuroD:Cre, NeuroD:IRES-GFP and either Cre inducible sh*Kif21b* #2 or sh-scramble. GFP-positive electroporated cells are depicted in green. Nuclei are stained with DAPI. Scale bars: (**C,E**) 50 μm.

**Figure S2. *Kif21b* depletion induces defects in tangential migration of interneurons, related to Figure 1.**

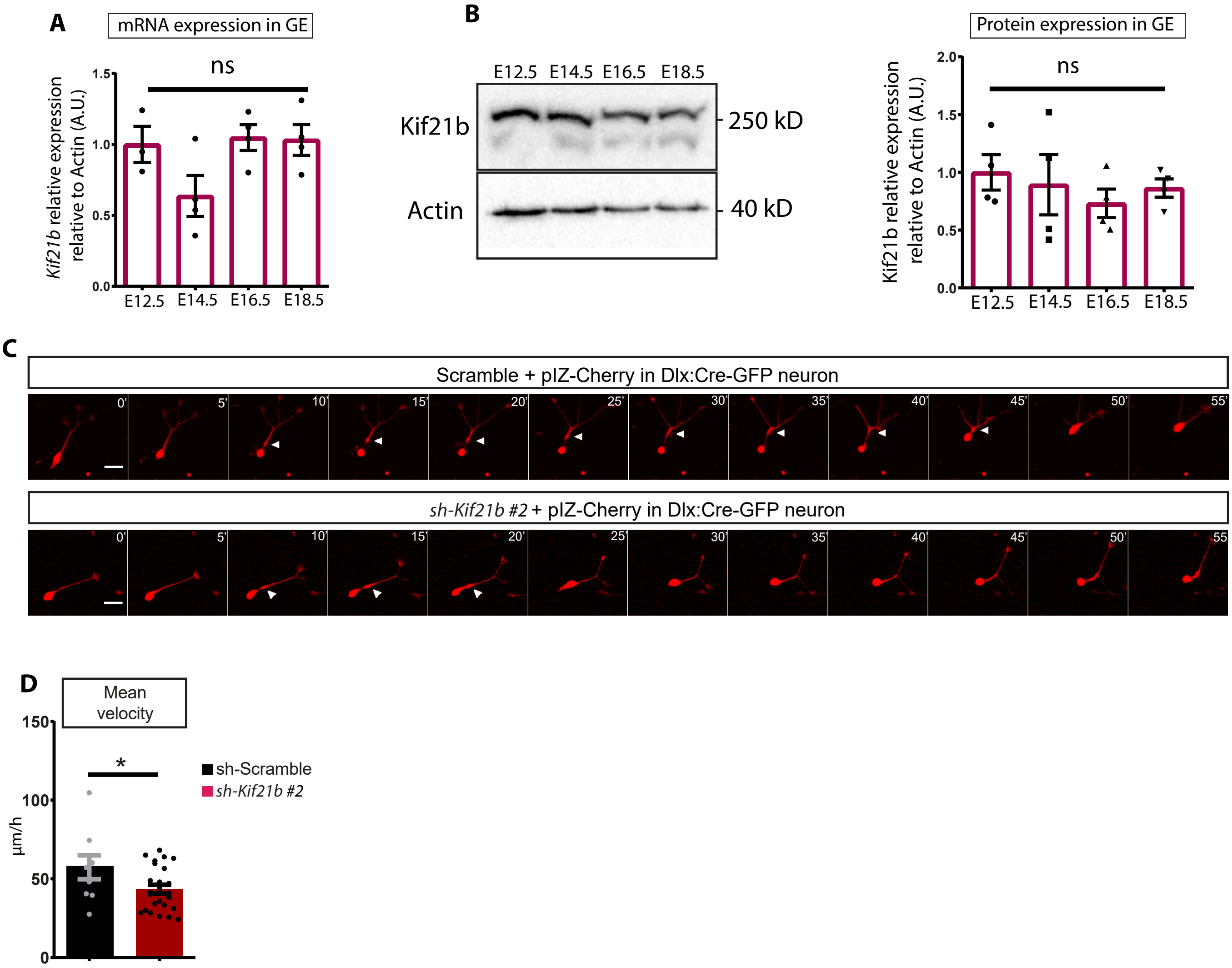


(**A,B**) RT-qPCR (n=4 brains per stage) (**A**) and Western blot (n=3 brains per stage) (**B**) analyses indicating expression of *Kif21b* transcripts and Kif21b proteins in mouse ganglionic eminences (GE) at different embryonic stages (from E12.5 to E18.5) (n=4 brains per stage). Data are represented as means ± s.e.m. Significance was calculated by one-way ANOVA, Bonferroni's multiple comparisons test, ns, non-significant. (**C**) Time-lapse sequence (min) showing the migration of interneurons out of MGE explants cultured from Dlx5,6 Cre-GFP E13.5 embryos, electroporated with a Cre-inducible cherry expressing plasmid together with Cre-inducible shRNA-*Kif21b* #2 or sh-scramble. Scale bar: 20 µm. (**D**) Quantification (means ± s.e.m.) of the mean velocity (µm/h) showing defects in tangential migration of *Kif21b*-depleed interneurons. Number of cells analyzed: scramble=9, sh-*Kif21b* #2= 25, from at least three embryos per condition in two independent experiments. Significance was calculated by unpaired two-tailed Student t-test, ns, non-significant; *P < 0.05.

**Figure S3. Overexpression of Kif21b-truncated constructs does not alter radial migration, related to Figure 3.**

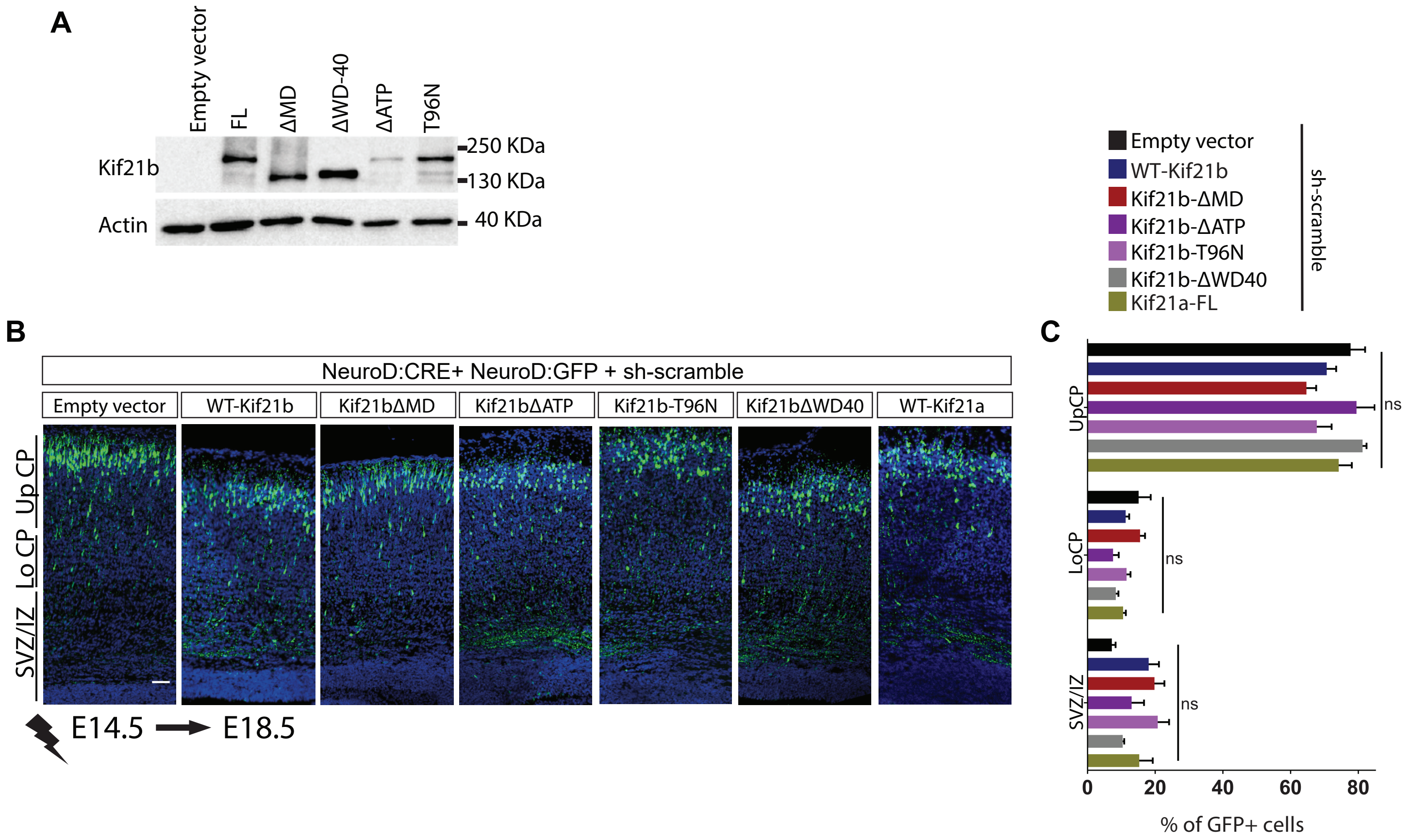


(**A**) Western blot of N2A cells transfected with the indicated full length (FL) or truncated Kif21b constructs expressed under the NeuroD promoter. Actin was used as a loading control. (**B**) Coronal sections of E18.5 mouse cortices electroporated at E14.5 with NeuroD:Cre and NeuroD:GFP together with sh-scramble and the different NeuroD:Kif21b constructs. GFP-positive electroporated cells are depicted in green. Nuclei are stained with DAPI. Scale bars, 50 µm. (**C**) Analysis (means ± s.e.m.) of the distribution of GFP-positive neurons in different regions (Up CP, Upper cortical plate; Lo CP, Lower cortical plate; SVZ / IZ, subventricular zone / intermediate zone) in all conditions as indicated. Number of embryos analyzed per condition: n≥3 for all conditions. Significance was calculated by two-way ANOVA, Bonferroni's multiple comparisons test. ns, non-significant; *$P < 0.05$.

**Figure S4. Kif21b interacts with actin and actin binding proteins and regulate actin dynamics, related to Figure 4.**

**A**

Table of main cytoskeleton interactors

| Protein | Gene name | Accession |
|---|---|---|
| Kinesin-like protein KIF21B | Kif21b | Q9QXL1 |
| Actin, cytoplasmic 2 | Actg1 | P63260 |
| Actin, cytoplasmic 1 | Actb | P60710 |
| Kinesin-like protein KIF21A | Kif21a | Q9QXL2 |
| Actin, alpha skeletal muscle | Acta1 | P68134 |
| Myosin regulatory light chain | Myl12b | Q3THE2 |
| Tubulin beta-6 chain | Tubb6 | Q922F4 |
| Drebrin | Dbn1 | Q9QXS6 |
| Unconventional myosin-Va | Myo5a | Q99104 |
| Coronin-1C | Coro1c | Q9WUM4 |
| Microtubule-associated protein 2 | Map2 | P20357 |
| Myosin light polypeptide 6 | Myl6 | Q60605 |

**B**

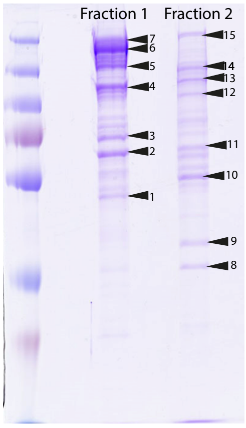


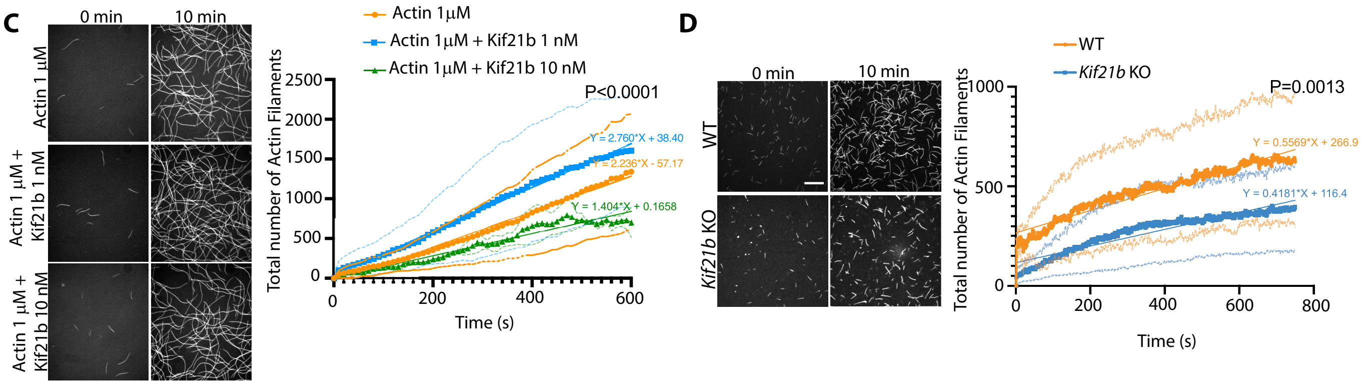


(**A**) List of cytoskeleton related partners found in the mass spectrometry analysis among the proteins that significantly interact with Kif21b in E18.5 cortices (see Table S1). (**B**) Isolated fractions of the recombinant Kif21b produced in BHK cells. The bands analyzed for contaminants are shown in arrowheads (see Table S2). (**C**) Representative spinning-disk images of *in vitro* analysis of actin nucleation (stained with phalloidin) at the beginning (0 min) and at the end (10 min) of the experiment in presence or absence of recombinant Kif21b (1 nM or 10 nM). Scale bar 10 µm. The graph shows the number of actin filaments assembled during a 600 s time-lapse. Number of replicates: Actin 1 µM, n=7; Actin 1 µM + Kif21b 1 nM, n=6; Actin 1 µM + Kif 10 nM, n=2. (**D**) *In vitro* analysis of nucleation of phalloidin-labeleld actin from protein extracts of WT or *Kif21b*$^{flox/flox}$ (KO) mouse cortices by spinning-disk microscopy. Scale bar: 10 µm. The graph shows the number of actin filaments assembled during a 600 s time-lapse. Three independent experiments were performed for both (WT and KO) conditions.
